# Promoter and Gene-Body RNA-Polymerase II co-exist in partial demixed condensates

**DOI:** 10.1101/2024.03.16.585180

**Authors:** Arya Changiarath, David Flores-Solis, Jasper J. Michels, Rosa Herrera Rodriguez, Sonya M. Hanson, Friederike Schmid, Markus Zweckstetter, Jan Padeken, Lukas S. Stelzl

## Abstract

In cells, transcription is tightly regulated on multiple layers. The condensation of the transcription machinery into distinct phases is hypothesised to spatio-temporally fine tune RNA polymerase II behaviour during two key stages, transcription initiation and the elongation of the nascent RNA transcripts. However, it has remained unclear whether these phases would mix when present at the same time or remain distinct chemical environments; either as multi-phase condensates or by forming entirely separate condensates. Here we combine particle-based multi-scale simulations and experiments in the model organism *C. elegans* to characterise the biophysical properties of RNA polymerase II condensates. Both simulations and the in vivo work describe a lower critical solution temperature (LCST) behaviour of RNA Polymerase II, with condensates dissolving at lower temperatures whereas higher temperatures promote condensate stability. Importantly this gradual change in temperature correlates with an incremental transcriptional response to temperature, but is largely uncoupled from the classical stress response. The LCST behaviour of CTD also highlights that these condensates are physio-chemically distinct from heterochromatin condensates. Expanding the simulations we model how the degree of phosphorylation of the disordered C-terminal domain of RNA polymerase II (CTD), which is characteristic for each step of transcription, controls demixing of CTD and pCTD in line with phase separation experiments. We show that the two phases putatively underpinning the initiation of transcription and transcription elongation constitute distinct chemical environments and are in agreement with RNA polymerase II condensates observed in *C. elegans* embryos by super resolution microscopy. Our analysis reveals how depending on its post-translational modifications and its interaction partners a single protein can adopt multiple morphologies and how partially engulfed condensates promote the selective recruitment of additional factors to the different phases.

## INTRODUCTION

Phase separation of biomolecules to form condensates provides a mechanism for cells to control processes in time and space. These condensates create local differences in protein concentrations, which is a way how phase separation can specifically promote or impede biological processes. Numerous proteins within cells have the potential to undergo phase separation under physiological conditions^1^. Should these proteins converge into a single location, condensation would neither increase local protein concentration nor enable specific biological processes. Specificity requires, therefore multiple co-existing phase-separated condensates to provide for spatio-temporal control over cellular processes^2^. As an additional layer of even finer regulation molecules may not only form distinct condensates that are entirely separated but also multiphasic condensates^3^. In multiphase condensates, there are distinct phases that do not mix. An essential question is what are the factors that determine the existence of multiple condensates and multiple phases in the condensates^4–6^. To understand the behaviour of condensates we have to describe three layers: the primary protein sequence, post transcriptional, reversible, chemical modifications, and how the two former alter and are altered by protein-protein interactions^7,8^.

To study the roles of multiphasic condensates, their co-existence, and how they might provide specific molecular recognition and thus cellular regulation, we turned to the C-terminal domain (CTD) of RNA Polymerase II (Pol II). As the central protein which transcribes genes from genomic DNA into mRNA the molecular mechanisms controlling RNA Pol II have been extensively studied over the last decades. Transcription (Fig 1A) is a linear process that can be divided into several regulatory steps, starting with recruitment of the Polymerase, an elongation process across the gene body, and finally transcriptional termination^9–11^. Recent experimental advances hypothesize that transcriptional initiation and post transcriptional processes associated with elongation might occur in different biomolecular condensates^10,12–17^. The formation of biomolecular condensates is frequently driven by multivalent interactions within intrinsically disordered regions of proteins and many proteins involved in transcription feature such disordered regions^18,19^. The CTD of RNA polymerase II is an intrinsically disordered protein domain^11^ that can form phase-separated condensates and its ability to form different condensates has been hypothesized to be important in the transcription of genes^13,20–22^ (Fig 1A). CTD forms condensates *in vitro* and contributes to RNA Polymerase II phase separation *in vivo*^13^. These condensates are thought to underpin the recruitment of RNA Pol II and the initiation of gene transcription, which are referred to as promoter condensates. CTD is then further phosphorylated as transcription initiation switches to transcription elongation and syntheses the RNA from the DNA template. Phosphorylated CTD forms condensate with proteins involved in transcription elongation and RNA processing^14,15^, which are called gene-body condensates^10^. Several proteins involved in regulating transcription elongation, such as P-TEFb directly or indirectly (splicing), have been observed to localise within these condensates. Whether such condensates will stay de-mixed or mixed when in direct contact remains unclear. Recently a surface condensation model was proposed to account for the dynamics of RNA polymerase II as it progresses from initiation and enters elongation as revealed by super resolution experiments in zebrafish^23^. However, splicing condensates remain de-mixed from initiation condensates^15^. To disentangle how condensates impact different stages of transcription it is therefore crucial to comprehensively characterise the phase behaviour of RNA Pol II and examine how phosphorylation affects condensates.

**Figure 1.**
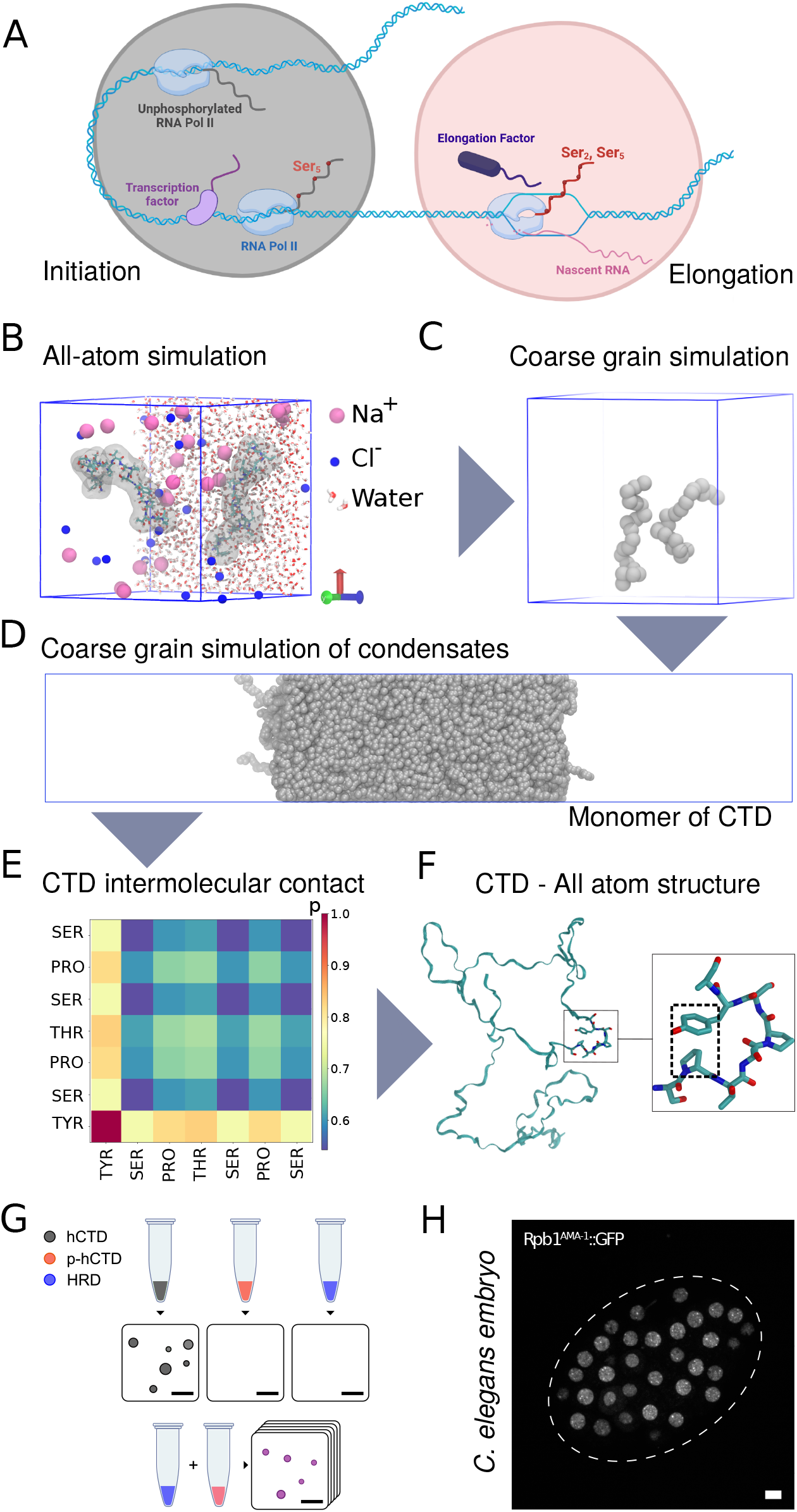
A) Hypothesized condensates underlying the initiation of gene transcription and the elongation phase. B) All-atom simulation capturing the behaviour of two 21mer CTD peptides. C) Coarse-grained(CG) simulation featuring two 21mer CTD peptides. D) CG slab simulation depicting CTD condensate. E) Interactions among CTDs in CG simulation revealed through intermolecular contact frequency. F) All-atom model of yeast CTD from Ref. 1 highlighting Tyr-Pro contacts. G) LLPS in vitro. Different combinations of hCTD, p-hCTD and HRD promote LLPS in vitro. H) In vivo imaging of RNA Pol II condensates in living embryos expressing GFP-tagged RPB1 (ama-1 in *C. elegans*).

Molecular simulations can serve as a “computational microscope”^24^ to elucidate drivers of cellular processes and phase behaviour on the molecular-scale^3,6,25–34^ and are especially powerful when combined with experiments^7,35^. Atomistic molecular dynamics (MD) simulations can resolve molecular interactions in condensates with high accuracy^7,36^. However simulating phase separation requires accessing long time and large length scales and thus the description of the system has to be simplified, by grouping atoms into particles, which is referred to as coarse-graining. Current coarse-grained models can capture important trends in how changes in sequence^37^ and post-translational modifications shape the single-chain and condensed phase behaviour of disordered proteins^27,38,39^. The loss of atomistic detail, begs the question whether all essential interactions are captured with sufficient accuracy. To ensure that simulations accurately describe phase behaviour requires a comparison of coarse-grained simulations to atomistic molecular dynamics and experiments reporting on biophysical properties^38^.

Here, we elucidate, using molecular dynamics, conditions under which the CTD, phosphorylation of the CTD (pCTD), and the binding of the HRD domain of cyclin T1 to pCTD, promote RNA Polymerase II condensate formation. Using the model organism *C. elegans*, we tested the predictions made by our simulations in a living organism under physiological conditions. Strikingly, in the simulations and *C. elegans* embryos, CTD forms condensate with lower critical solution temperature (LCST) behaviour. This clearly distinguishes it from other previously characterised biological condensates in the nucleus, which typically dissolve at high temperatures, and hints at discrete chemical properties intrinsic to IDRs that could contribute to the co-existence of multiple distinct condensates in the nucleus. Our simulations show that CTD and pCTD-HRD indeed do not mix but instead form (partially) demixed phases, in line with experiments with purified CTD, pCTD and HRD. The degree of demixing is determined by the extent of phosphorylation. In simulation with hyperphopshorylated pCTD, the pCTD-HRD phase engulfs the CTD phase, which we demonstrate by developing a general simulations approach to predict the morphologies of condensates. The simulations further predict that the exact degree of engulfment would vary based on conditions such as condensate composition. To ensure that our simulations accurately capture the molecular driving forces of CTD phase separation, we compare coarse-grained simulations to atomistic molecular dynamics, nuclear magnetic resonance (NMR) based structural ensembles. In agreement with simulations, we find in super-resolution microscopy that in *C. elegans* embryos CTD is indeed fully and partially engulfed by pCTD. While the two condensates are largely de-mixed and chemically distinct, they remain in close contact, which may underpin the efficient movement of RNA Polymerase II from initiation to elongation as part of the functional cycle of RNA Polymerase II.

## METHODS

### Coarse-grained simulation model

We employed coarse-grained molecular dynamics simulations using the residue-level hydropathy scale (HPS) model which represents amino acids as beads and the solvent effect is considered implicitly^25^. The HPS model intricately captures the physical and chemical attributes inherent to the natural ensemble of 20 amino acids into these beads, resulting in sequence-specific interactions among them. Three kinds of interaction are considered between the beads. Firstly, beads are bonded to each other through harmonic potential energy, *V*_*H*_ (*r*) = *k*(*r* – *r*_0_)^2^/2 where 10 kJ/ Å^2^ is force constant and *r*_0_ = 3.8 Å is the bond length. Secondly, electrostatic interaction between charged amino acids is modelled using the Debye–Hückel (DH) electrostatic screening term.

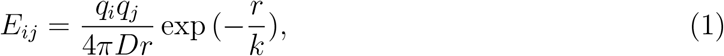

where *k* = 1 nm is the Debye screening length which corresponds to a salt concentration of 100 mM and *D* = 80 is the dielectric constant of the aqueous solvent. A cut-off distance of *r*_*cut*_ = 3.5Å is set. If *r* is greater than *r*_*cut*_, then *E*_*ij*_ is set to zero. Lastly, the short-range pairwise interaction between beads is represented by Ashbaugh–Hatch potential,

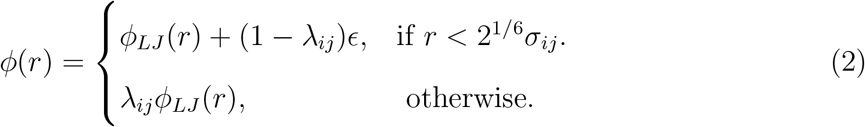

where *ϕ*_*LJ*_ is the Lennard-Jones potential,

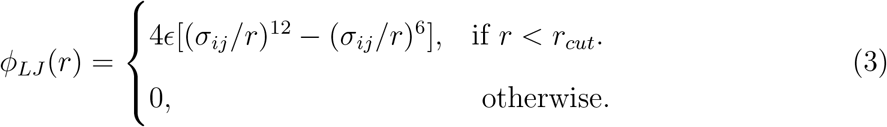

and *λ*_*ij*_ and *σ*_*ij*_ are the arithmetic average of the hydrophobicity scale and size of amino acids, respectively. A cut-off distance of *r*_*cut*_ = 2.00Å is used and the potential is shifted accordingly. The residue specific hydrophobicity values range from 0 to 1 as the hydrophobicity increases. The HPS model also provides parameters for phosphorylated amino acids such as phospho-serine^8^, which have been used to model the effect of phosphorylation on the fifth position in the CTD heptad repeat motifs, which we refer to as Ser_5_. To study temperate-dependent phase behaviour we used the HPS-T model, a version of the HPS model which has been extended to predict the temperature dependent properties of disordered proteins^40^.

Coarse-grained molecular dynamics simulations were run with the HOOMD-Blue package^41^. The system is initialized by placing the protein chains inside an extended rectangular box with a minimum distance between protein chains. The simulations were performed under *NVT* ensemble conditions, using a Langevin thermostat with a friction coefficient of 0.01*ps*^−1^. The proteins are equilibrated within the first few nanoseconds by condensing into a liquid dense and dilute phase. The density profile is extracted from the protein concentration along the *z* axis as described by Tesei et al.^27^.

### Atomistic molecular dynamics simulations

Two 21mer CTD fragments were simulated in all-atom molecular dynamics with explicit solvent. We employed the Amber99sb-star-ildn-q protein force field^42–45^ and the TIP4P-D water model^46^ in the simulations. Equations of motions were integrated with a 2 fs time step. The *NPT* ensemble was maintained at 300 K and 1 bar with the Bussi-Donadio-Parrinello thermostat^47^ and the Parrinello-Rahman barostat^48^. Simulations were run in GROMACS^49^, with a cumulative simulation duration of 39.8 µs.

### Simulated systems

Employing coarse-grained molecular dynamics simulations enables us to gain an in-depth understanding of the complexities involved in the regulation of gene transcription. First, we simulated CTD proteins from *C*.*elegans, Saccharomyces cerevisiae, Drosophila melanogaster*, and *Homo sapiens* (Fig S2). To understand how deviations from ideal heptad repeat sequence Y_1_S_2_P_3_T_4_S_5_P_6_S_7_ affect phase behaviour we also conducted simulations of chains of the same length as these natural proteins, but with idealized sequences. Simulations of *C. elegans* CTD were also run with the HPS-T model to investigate its temperature dependent phase behaviour^40^.

To study the formation of distinct phases or multiphasic condensates we focus on a simplified system, comprising three key components that play a vital role in the transcription process and may underlie key aspects of phase behaviour in the nucleus. These elements consist of CTD of RNA polymerase II (140 mer with 20 ideal heptad repeats), responsible for the promoter condensate; phosphorylated CTD (140 mer with various degrees of phosphorylation); and Histidine Rich Domains (HRD) of P-TEFb Cyclin T1, contributing to the formation of the gene body condensate (Fig. 1A).

Analysing the properties of these condensates provides insights into their role in gene regulation. Additionally, the coarse-grained nature of the simulation allows us to explore large-scale and extended-time thermodynamic and mechanical properties, crucial for understanding the stability and structure of multiphasic condensates. We initiated our study by examining the formation and regulation of the promoter condensate, which is formed through the phase separation of CTD. We explored the impact of phosphorylation on CTD phase separation and the co-phase behaviour of HRD with CTD and pCTD through a series of simulations. The coacervation behaviour of HRD and pCTD was explored using the flory-huggins theory. To gain a comprehensive understanding of how phosphorylation impacts the formation of multiphase condensates, we conducted simulations involving CTD, pCTD, and HRD, systematically varying the phosphorylation levels of the pCTD chains in each simulation. Furthermore, to explore the structure and stability of these multiphase condensates, we carried out simulations of droplets of these condensates in a bigger cubic box.

### Analysis of phase behaviour with a mean-field model

We analyse the simulated phase diagram of pCTD and HRD co-phase separation with a mean-field Flory-Huggins model^50^,

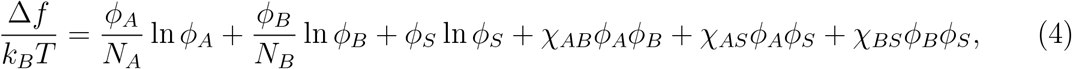

Here, *ϕ*_*i*_ and *N*_*i*_ represent the volume fractions and relative degrees of polymerisation and *χ*_*ij*_ the binary interaction parameters. In the model, *ϕ*_*S*_ follows from the assumption of incompressibility: *ϕ*_*S*_ = 1 − *ϕ*_*A*_ − *ϕ*_*B*_. Subscripts *A, B*, and *S* respectively denote HRD, pCDT, and “solvent”. The relative sizes of the polymers, which we fix at *N*_*A*_ = 70 and *N*_*B*_ = 140, have been normalized by a scaled monomeric volume (*N*_*S*_ = 1).

The mean-field phase diagram was obtained in the usual way by demanding that the chemical potential and the osmotic pressure be similar in the dilute and dense phase, using well known methodologies^51^.

### Calculation of interfacial tension

Interfacial tension is determined using two methods employing capillary wave theory: first, by assessing interfacial width in slab simulations, and second, by analysing shape fluctuations of droplets.

In slab geometries, capillary wave theory (CWT) with the block analysis method was employed to estimate the interfacial tension of condensates^52^. In the block analysis method, the interface region is divided into rectangular segments with dimensions of *L*_*x*_*/n*_*B*_ × *L*_*y*_*/n*_*B*_ × *L*_*z*_, where *n*_*B*_ is an integer, where we choose 3 < *n*_*B*_ < 8. Within each segment, intrinsic profiles are computed based on the density function:

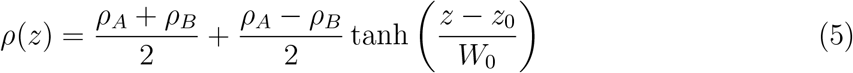

Where *ρ*_*A*_ and *ρ*_*B*_ represent the bulk densities of two phases, denoted as *A* and *B*, respectively. *z*_0_ is the position of the interface, while *W*_0_ corresponds to the intrinsic width of the system. The apparent interfacial width (*W*) at a finite temperature is determined by combining the intrinsic width *W*_0_ and the capillary contribution:

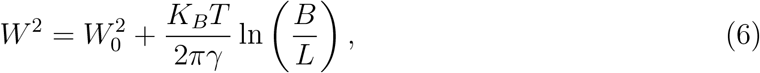

where *k*_*B*_*T* is the thermal energy, *γ* represents the interfacial tension, and *B/L* is the ratio of block size to system size. The apparent interfacial width (*W*) is computed as a function of the block size and interfacial tension can be calculated using equation 6. With CWT, we can determine the interfacial tension at the interface between a dense protein phase with the solvent, as well as at the interface between two distinct protein phases.

We compared the interfacial tension of the homogeneous condensate, as obtained from the above calculation, with the interfacial tension obtained using Kirkwood-Buff relation^53^,

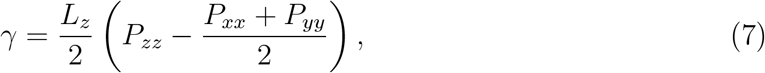

where, *P*_*αβ*_ are the components of the pressure tensor.

Furthermore, the interfacial tension of a droplet can be estimated from the thermal fluctuations of its spherical shape based on the method pioneered by Henderson and Lekner^54^ which has found recent application in the analysis of biological condensates^26^. The potential energy (*U*) associated with these shape fluctuations is primarily determined by the interfacial tension (*γ*), which can be mathematically expressed as *U* = *γδA*. Here, *δA* is the small variation in interfacial area relative to that of a perfect sphere. By analysing the droplet’s shape fluctuations at the lowest order using a spherical harmonics expansion, we can derive two independent measures of the interfacial tension:

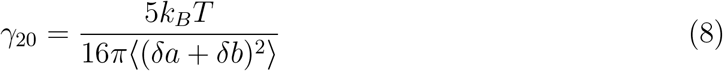

and

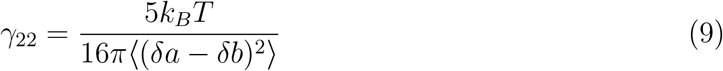

Here, the droplet’s instantaneous shape is described by a general ellipsoid, with *δa* and *δb* representing the lengths of any pair of principal axes of the ellipsoid, and *R* indicating the average radius of the droplet. By selecting the relevant protein chains, we computed surface tensions of the homogeneous protein phases as well as interfacial tensions between different protein phases.

### Characterisation of disordered protein conformation

In our study, we investigate the impact of phosphorylation on the conformation and compactness of CTD chains in dilute and dense phases. A key metric for this analysis is the radius of gyration (*R*_*G*_). Additionally, we assess protein anisotropy to identify structural changes resulting from phosphorylation. The gyration radius of chains is calculated using the following equation:

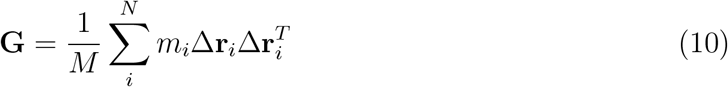

Here, M represents the mass of the chain, *N* is the number of monomers in the chain, andΔ**r**_*i*_ is the position vector of monomer *i* with respect to the chain’s center-of-mass. The root mean square radius of gyration is calculated from the diagonal elements of the **G** matrix, given by *R*_*g*_ = ⟨*Gxx* + *G*_*yy*_ + *G*_*zz*_⟩^1/2^. Furthermore, we examined the anisotropy of the chain using the following equation^53^:

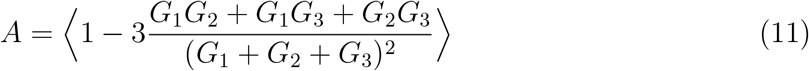

Here, *G*_1_, *G*_2_, *G*_3_ represent the three eigenvalues of the matrix **G**. Analysing the anisotropy of the chain allows us to characterise any structural modifications and deviations from a symmetric conformation that may arise due to phosphorylation.

### Interactions of disordered protein chains in condensates

The excess of monomer A in the vicinity of a B monomer, 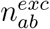 is calculated using the equation:

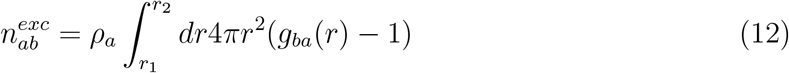

Here, *r*_1_ and *r*_2_ represent the minimum and maximum radial distances from the B monomer, respectively, over which the excess number of monomer A is computed. The specified cutoff values are *r*_1_ = 0.45, nm and *r*_2_ = 7.5, nm. The correlation function *g*_*ba*_(*r*) is defined as:

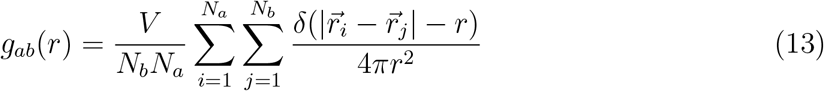

Here, *N*_*a*_ and *N*_*b*_ represent the total number of monomers of type A and B, respectively. The excess of monomer, 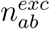, functions as a metric for quantifying the degree of protein-protein interactions.

### Protein expression and purification

Plasmids were adapted to produce the carboxyl-terminal domain of human Pol II (hCTD; RPB1 residues 1593–1970) and the Histidine Rich Domain (HRD; Cyclin-T1 residues 195-265). The constructs are composed of histidine (6xHis) and maltose-binding protein (MBP) tags located at the N-terminus. A flexible linker of ten consecutive asparagines and the tobacco etch virus (TEV) protease cleavage site were introduced to allow cleavage of the tags. The protein sequences were codon-optimized for expression in bacteria. For site-specific labeling with a fluorescent tag, a cysteine residue was present at the N-terminus downstream of the TEV cleavage. MBP-tagged proteins were overexpressed in *E. coli* BL21 RP-Codon Plus DE3 cells (hCTD; Agilent Cat. 230255) and BL21 DE3 (HRD; Novagen-Merck) at 37 °C in LB media. Cells were collected after overexpression (10 min, 10000 x g; Avanti JXN-26, Beckman Coulter) and resuspended in lysis buffer (25 mM HEPES, pH 7.4, 300 mM NaCl, 30 mM imidazole, cOmplete EDTA-free protease-inhibitor cocktail, 0.1 mg/l lysozyme and 0.1 mM PMSF) at 4 °C. Cells were disrupted by sonication (15 s pulse at 60 W, 45 s pause, 10 min total; SONOPULS, Bandelin). The cell extract was clarified by centrifugation (20 min, 45000 x g; Avanti JXN-26, Beckman Coulter), loaded with a sample pump (Äkta Pure GE Healthcare) into an ion-metal affinity chromatography column (IMAC; FastFlow-Hitrap GE Healthcare), and eluted with imidazole. The purity of the fractions was improved by size exclusion chromatography (Superdex 75 26/600 GE Healthcare) and Anion-Exchange Chromatography (MonoQ 10/100 GL GE Healthcare). Fractions containing the MBP-tagged protein were merged and concentrated. TEV protease was added (1:100 mass ratio) to the mix for cleavage. The reaction was incubated overnight (16-18 hours) at 4 °C with gentle agitation. Cut tags were removed using IMAC purification, collecting and concentrating the unretained fractions. For the phosphorylated version of hCTD, mixtures were incubated overnight (37 °C) MBP-hCTD and CDK9/CyclinK (Promega; 1000:1 mass ratio) in HEPES 25 mM; NaCl 150 mM; MgCl2 5 mM; ATP 10 mM and DTT 1 mM at pH 7.4 to achieve maximum phosphorylation. A Small portion of MBP-p-hCTD sample was cut as described above, purified by Anion-Exchange Chromatography (MonoQ 5/50 GL GE Healthcare) and analyzed by mass spectrometry to estimate de degree of phosphorylation. Fast protein liquid chromatography (FPLC) was performed at 4 °C using an Äkta Pure system (GE Healthcare). HRD purified protein was collected from Reversed-Phase HPLC (preparative column: Vydac 214TP 5 µm C4, 250 x 10mm; A: water + 0.1% TFA, B: acetonitrile + 0.1% TFA; HPLC system JASCO with diode array detector) and the molecular weights were confirmed by mass spectrometry (analytical column: Waters BioResolve RP mAb, Polyphenyl, 450A, 2,7 m, 4.6 x 100mm; A: water + 0.1% TFA, B: acetonitrile + 0.1%TFA; LC-MS: Acquity Arc System, Waters, with SQD2-Mass-Detector: Single Quadrupole; Direct mass: ZQ 4000 Waters, Single Quadrupole, injection by syringe pump). HPLC samples were lyophilized for further experiments.

### A. In vitro phase separation

Stock solutions for MBP-hCTD and MBP-p-hCTD (200 µM) were measured by nanodrop (Thermo) and weighting dry protein (HRD; 1 mM) and dissolving it in pre-cooled buffers at 4 °C. Buffer solutions were filtered (0.22 µm) after preparation. Dextran T400 (Pharma) was used as a crowding agent at 5 % w/v. MBP-hCTD and MBP-p-hCTD were labeled with Alexa Fluor 488 and 594 maleimide, respectively (AF488; AF594) following the protocol in the microscale kit (Invitrogen). Micromolar amounts (¡ 1.5 µM) of fluorescently labeled protein were mixed with unlabeled protein to reach the final concentrations. Prior to imaging, samples were gently mixed by pipetting. Five microliters of sample were loaded onto glass slides and covered with ø18 mm coverslips. Differential interference contrast (DIC) and fluorescence micrographs were acquired at room temperature using a Leica microscope (DM6000B) equipped with a 63x/1.20 objective (water immersion) in 25 mM HEPES, pH 7.4, 150 mM NaCl and 5 % w/v Dextran. Micrographs were analyzed and processed with Fiji (NIH). Micrographs are representative of at least 3 independent replicates

### *C. elegans* strains and maintenance

AMA-1::GFP (ama-1(ot1037[gfp::ama-1]) IV.) strains were provided by the CGC, which is funded by NIH Office of Research Infrastructure Programs (P40 OD010440). Unless otherwise indicated, experiments were performed using early embryos isolated from animals cultured at 20 °C.

### Live microscopy and Immunofluorescence (IF)

Fluorescent images of live animals or embryos were captured in M9 buffer on 2% agarose pads, embryos were incubated at the indicated temperatures for 1h prior to imaging. Images of live embryos were captured using the LAS X Life Science Microscope Software with the Stellaris 8 FALCON (Leica) equipped with an HC PL APO 100x/1.40 OIL STED WHITE objective lens and a white light laser source. Z-stacks were recorded with a z-spacing of 200 nm and an xy-pixel size of 54 nm. All images were deconvolved using the inbuild LIGHTNING deconvolution module (Leica). For IF, bleached embryos were fixed for 5 min in 1% formaldehyde before spotting onto poly-l-lysine–coated coverslips, freezing on dry ice, and storing at −80 °C. When needed, slides were freeze cracked and immediately fixed in a −20 °C 100% ethanol (EtOH) bath for 2 min and then dried. After washing 3 × 5 min with PBS +0.25% Triton X-100 (PBS-T) and blocking for 1 h with PBS-T and 2% milk, slides were incubated overnight at 4°C in a humid chamber with primary antibodies in PBS-T and 2% milk: 1:500 mouse anti-H3K9me2 MABI0317 (MBL) and 1:500 polyclonal rabbit anti-RFP (Rockland). After washing 3 × 5 min with PBS-T, slides were incubated for 1 h at RT in a humid chamber with secondary antibodies in PBS-T and 2% milk: 1:1,000 goat anti-mouse Alexa Fluor 488 (A11001; Invitrogen) and 1:1,000 donkey anti-rabbit Alexa Fluor 555 (A31572; Invitrogen). DNA was counterstained with DAPI (1:2,000) in PBS-T for 10 min, washed 3 × 5 min with PBS-T, and mounted with ProLong Gold Antifade (Thermo Fisher Scientific). SIM images were acquired using an ELYRA 7 LS mit 2x pco.EDGE 4.2 (Zeiss) and a Plan-Apochromat 100x/1,4 Oil objective lens.

We quantified nuclear foci using the KNIME Analytics Platform^55^. In summary, nuclei were detected using a seeded watershed segmentation using the DAPI channel (SIM images) or the green channel. For foci detection, we used a Laplacian-of-Gaussian detector from TrackMate^56^ (fmi-ij2-plugins-0.2.5, https://doi.org/10.5281/zenodo.1173536) on the AMA-1 (GFP) channel. Potential foci outside of a nucleus were ignored in the analysis. Results were plotted per condition using the R package ggplot2 (https://ggplot2.tidyverse.org/).

### RNA sequencing

RNA was extracted from early embryos from a synchronized culture grown at 20°C and exposed to indicated temperatures for 2 h, using Trizol as previously described 6. Embryos were freeze cracked five times, then RNA was extracted with chloroform followed by isopropanol precipitation. Further purification was performed with the RNA Clean and Concentrator kit (Zymo). Libraries were produced using the Stranded Total RNA Prep Ligation with Ribo-Zero Plus Kit (Illumina). rRNA was depleted using the Ribo-Zero Plus kit supplemented with custom made *C. elegans* specific rDNA oligos (IDT). Libraries were profiled in a DNA 1000 Chip on a 2100 Bioanalyser (Agilent technologies) and quantified using the Qubit 1x dsDNA HS Assay Kit, in a Qubit 4.0 Fluorometer (Life technologies). Equimolar amounts of indexed libraries were pooled and sequenced on a NextSeq 2000 (Illumina). Reads were analysed as described previously^57^. Adapters were trimmed using Trimmomatic v0.39. Reads were aligned to the *C. elegans* genome (ce10) with the R package QuasR v1.42.1, (www.bioconductor.org/packages/2.12/bioc/html/QuasR.html). The command `“proj <-qAlign(“samples.txt”,”BSgenome. Celegans.UCSC.ce10”, splicedAlignment=TRUE)”` instructs hisat296 to align using default parameters, considering unique reads for genes and genome wide distribution. Count Tables of reads mapping within annotated exons in WormBase (WS220) were constructed using the qCount function of the QuasR package to quantify the number of reads in each window `(qCount(proj, GRange_object,orientation=“same”))` and normalized by division by the total number of reads in each library and multiplied by the average library size. Transformation into log2 space was performed after the addition of a pseudocount of 8 to minimize large changes in abundance fold change (FC) caused by low count numbers. The EdgeR package v4.0.14 was applied to select genes with differential transcript abundances between indicated genotypes (contrasts) based on false discovery rates (FDR) for genes. Replica correlations are shown in Fig S5A of GO-Terms were extracted from wormbase.org and the GO-Term and KEGG pathway enrichment analysis was performed using the gprofiler2 package (v 0.2.2) as an interface to g: Profiler^58^.

## RESULTS

### Molecular drivers of CTD phase separation and functional implications

We first established that residue-level coarse-grained molecular dynamics simulations^25^ are a suitable model of the interactions of RNA pol II CTD (Fig 1B-F) and in turn, simulations revealed how phase separation is driven by the interactions of the heptad repeats and how deviations from the ideal heptad sequence and truncation affect the tendency of CTD to phase separate. To ensure that our simulations capture the essential elements of the behaviour of CTD, we compare to experiments and atomistic reference simulations reporting on the contact statistics of CTD at the level of condensates, inter-chain interactions, and intra-chain interactions (Fig 1B-F and Fig S1), which sets the scene of investigating functional aspects of CTD. In coarse-grained simulations, we find that the CTD (140mer with 20 ideal heptad repeats) phase separates on its own. Contacts between aromatic Tyr residues within the CTD are particularly prominent, as demonstrated in contact analysis of CTD (Fig 1E). These observations are consistent with previous experimental evidence highlighting the crucial role of hydrophobic interactions in driving phase separation^13,22^.

Tyr-Try contacts are prominent in the interactions of two 21mer CTD fragments in extensive atomistic reference simulations (39.8 µs in total) (Fig S1 C). By comparison of these atomistic reference simulations (Fig S1 A, B Movie S1) of two CTD fragments to coarsegrained simulations, we could also show that our coarse-grained molecular dynamics simulations capture CTD interactions not only qualitatively, but also to the overall magnitude of interactions as expected from atomistic molecular dynamics simulations (Fig 1B, C). The affinity for CTD fragments for each other is estimated, assuming a kinetic rate model^59,60^, to be *K*_*d*_ = 2.76 ± 0.2 mmol and *K*_*d*_ = 1.19 ± .005 mM for the atomistic and the coarse-grained simulations respectively (SI Text).

The contact analysis also highlights Tyr-Pro contacts (Fig 1E) which were recently highlighted by nuclear magnetic resonance (NMR) experiments and atomistic modelling to generate ensembles of all-atom CTD structures of human and yeast CTD, which account for the experimental data (Fig 1F)^22^. Finally, we compare simulations to experiments with CTD truncations, which shows that our modelling captures functional relevant trends in the behaviour of CTD. As anticipated, shorter CTD sequences exhibited less phase separation than longer ones^13^ (Fig S4). Additionally, we observed that the actual sequences from *Drosophilia melanogaster, C. elegans, Saccharomyces cerevisiae*, and *Homo sapiens*, which all feature deviation from the heptad repeat pattern Y_1_S_2_P_3_T_4_S_5_P_6_S_7_, to be less prone to phase separate than sequences of equal length which feature only copies of the ideal heptad repeat. In line with experiments, a truncated version of *Drosophilia* (20 repeats) CTD^61^, which features only idealised CTD repeats phase separates about as much as full-length *Drosophilia* real CTD, whereas the full length ideal *Drosophilia* CTD showed a considerably higher tendency to phase separate (Fig S4).

### CTD condensates dissolve at lower temperatures but are stable at room temperature in simulations and *C. elegans* unlike other nuclear condensates

Phase separated condensates are highly temperature dependent^40^ and indeed measuring temperature dependent changes in condensate behaviour *in vivo* is one of the gold standards in the field^62^. To test our model, we therefore calculated the temperature dependent phase behaviour of RNA Pol II CTD (Fig 2A-D, S2). We then compared the predicted temperature dependent changes in condensate behaviour to the behaviour of RNA Pol II condensates in *C. elegans* embryos, exposed to different temperatures. To quantify RNA Pol II condensates *in vivo* we imaged living embryos expressing the main RNA Pol II subunit RPB1 (ama-1 in *C. elegans*) tagged with GFP. Worms were cultured at 20°C (standard laboratory condition) and exposed for 1 h to a range of temperatures (4°C-30°C) (Fig 2E). Using automated image analysis we segmented RPB1::GFP condensates and quantified condensate number, size, and fluorescence intensity (Fig 2F-H). Strikingly we observed a high agreement between the simulations and the *in vivo* behaviour of RPB1 condensates. Interestingly, in contrast to most other described condensates which dissolve at higher temperatures and display an upper critical solution temperature (UCST), both the simulation model and the *in vivo* condensates display a lower critical solution temperature (LCST), resulting in reduced condensate formation at low temperatures and an increase in condensate formation at higher temperatures. Such LCST behaviour has recently been detected for RNA Pol II CTD in vitro^22^, which is in line with our *in vivo* experiments and simulations. LCST behaviour has been thoroughly investigated for synthetic polymers and more recently for disordered proteins^63–67^. Many Pro-rich protein sequences show LCST behaviour. Water molecules are highly organised around Pro residues^68^ and the formation of such water networks is associated with an entropic cost. At higher temperatures, the water network around Pro residues is less favourable and Pro is more prone to interact with other residues, favouring compact chain conformations and protein condensate formation. Other nuclear condensates, e.g., the heterochromatin condensates formed by the histone methyl transferase MET-2^57^ and HP1*α*^69^ dissolve with increasing temperatures and these differences in their response to temperature suggest that multiple distinct condensates co-exist in the nucleus.

**Figure 2.**
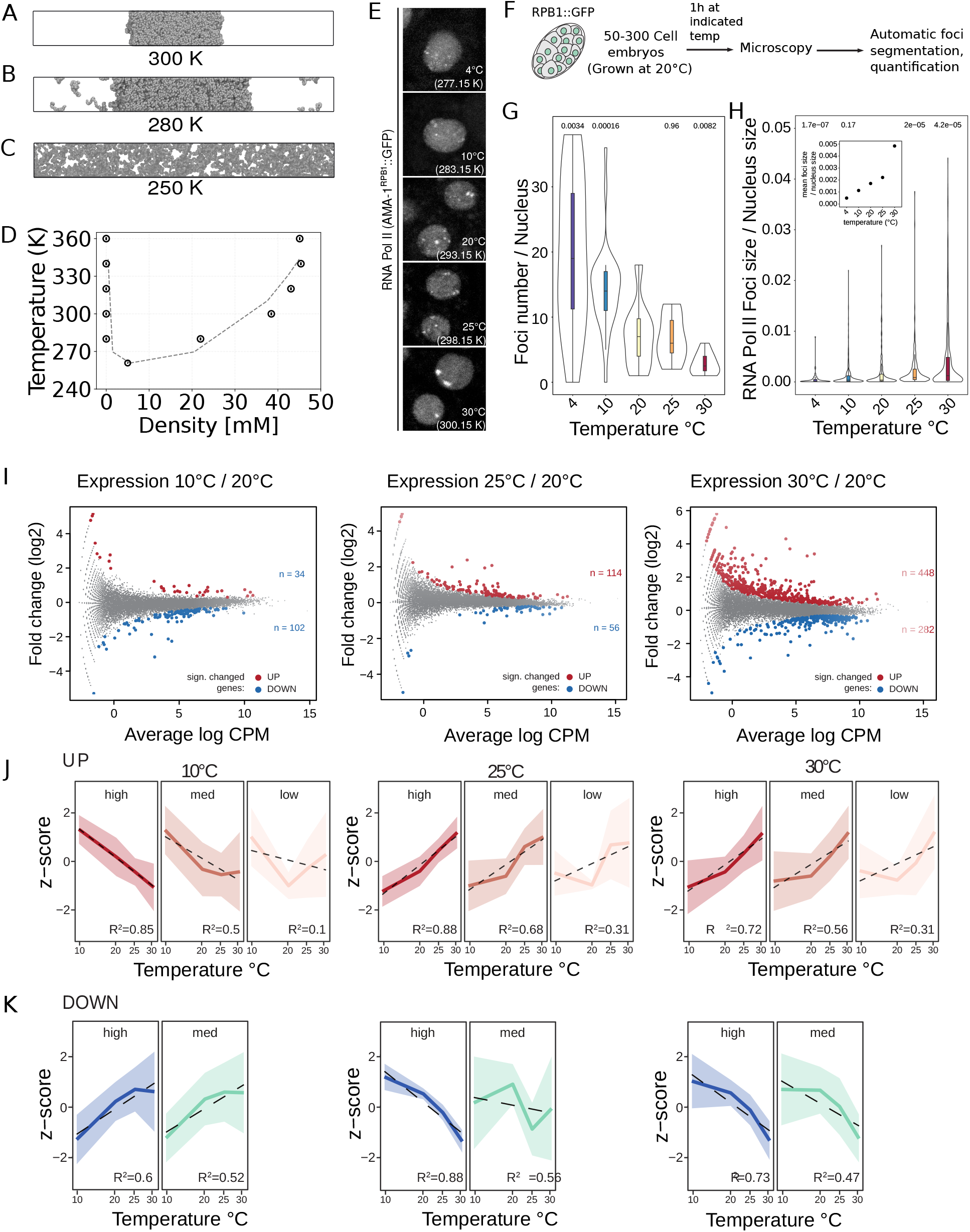
Temperature dependent phase behaviour of RNA Pol II CTD. A-C) Snapshots from coarse-grained simulations at 250 K, 280 K and 300 K. The system phase separates at higher temperatures. D) Phase diagram illustrating LCST phase behaviour. The uncertainties are indicated. E) RNA Pol II foci in *C. elegans* embryos at different temperatures. As the temperature rises, the number of foci decreases, while their size increases. F) Experimental setup for investigating the effect of temperature on CTD condensate in *C. elegans* embryos. G) The number of foci per nucleus. H) The size of foci relative to the total nuclear volume. Exposing *C. elegans* embryos to a range of temperatures from 10°C-30°C results in a gradual transcriptional response. I) Mean expression (CPM) versus expression change (log2 fold change) plots comparing transcriptome of embryos cultured at 20*C* (standard laboratory conditions) to embryos exposed to 10°C, 25°C, or 30°C for 2hrs. n indicates the number of deferentially expressed genes (*FDR* < 0.05), highlighted red significantly unregulated genes (fold change > 2), and blue significantly downregulated genes (fold change < −2) in all three plots.(Caption continues next page.) J) Expression changes of significantly up and downregulated genes identified in (a) at each temperature plotted as the mean z-score (solid line) and its standard deviation (shaded area). Dotted black line indicates a fitted linear regression. Each gene set was divided according to its mean expression level at 20°C (high> 75 percentile; med< 75 and > 25 percentile; low< 25 percentile).

### Temperature dependent changes in embryonic gene expression are largely different from the classical stress response

The impact of the temperature shifts on the *in vivo* RNA Polymerase II condensates raises the question whether, or how these gradual changes in temperature affect the embryonic transcriptome. We therefore sequenced total RNA from embryos exposed to 10°C, 20°C, 25°C and 30°C for 2 h. When compared to embryos cultured at 20°C (standard laboratory conditions) we observed a modest downregulation of genes at 10°C, while exposing worms to 25°C resulted in a predominantly upregulation of genes, which was further pronounced at 30°C (Fig 2I, S5). Interestingly these transcriptional changes showed a linear correlation with temperature (Fig 2J,K). This is in contrast to what would be expected from a stress response where a dedicated signaling cascade triggers a defined transcriptional program upon reaching an activation threshold^57^. Indeed none of the temperatures result in an up- or downregulation of genes associated with known stress response pathways comparable with the effect that a 1 h heat shock at 37°C has. We note that there are two notable exceptions hsp-70 and hsp-16.41. Both of these are heatshock proteins that show a gradual increase in expression from 10°C-30°C (Fig S3). Instead, the gene ontology terms (GO-Terms) enriched in the changed genes indicate that these small changes in temperature have a gradual impact on the expression of genes involved in metabolic processes. While it is a correlation, the gradual, linear change in transcription suggests that the biophysical properties of RNA Polymerase II may play a direct role in fine-tuning transcription.

### Co-condensation of Ser_5_ phosphorylated pCTD is driven by valence but not pattering

In addition to the separation of active from inactive regions in the genome, transcription itself occurs in multiple defined steps. The post-transcriptional phosphorylation state of RNA Polymerase II CTD is characteristic of each of these transcriptional stages^9,11^ and has been implicated in condensate formation. Here we ask how the degree of CTD phosphorylation and the distribution of phosphosites impacts condensate formation. To specifically examine how phosphorylation influences phase separation, we conducted simulations using CTD and pCTD chains, each comprising 140mers (20 repeats of the ideal heptad repeats)(Fig 3C, S7 A,D). We incrementally phosphorylated 50 CTDs in our simulations and in accordance with experiments, we find phosphorylation dissolves CTD condensates in a dose-dependent manner. Phosphorylating up to 15 Ser_5_ in the heptad motif out of a total of 20 Ser5 in the 140mer pCTD, still predominantly leads to partitioning into the dense phase of CTD. However, as the phosphorylation level of pCTD increases above the threshold of 15 Ser_5_, the dilute concentration of pCTD rises gradually as the electrostatic repulsion between pCTDs in the dense phase dominates and expels the pCTD chains into the dilute phase (Fig S7 B). The contact map illustrating pCTD-CTD interactions (Fig S6 B) highlights the relatively weak nature of the intermolecular binding between the CTD and pCTD with all the 20 Ser_5_ phosphorylated, as compared to the intermolecular interactions observed between CTD-CTD. pCTD with each heptad repeat phosphorylated does not phase separate on its own (Fig 3B).

**Figure 3.**
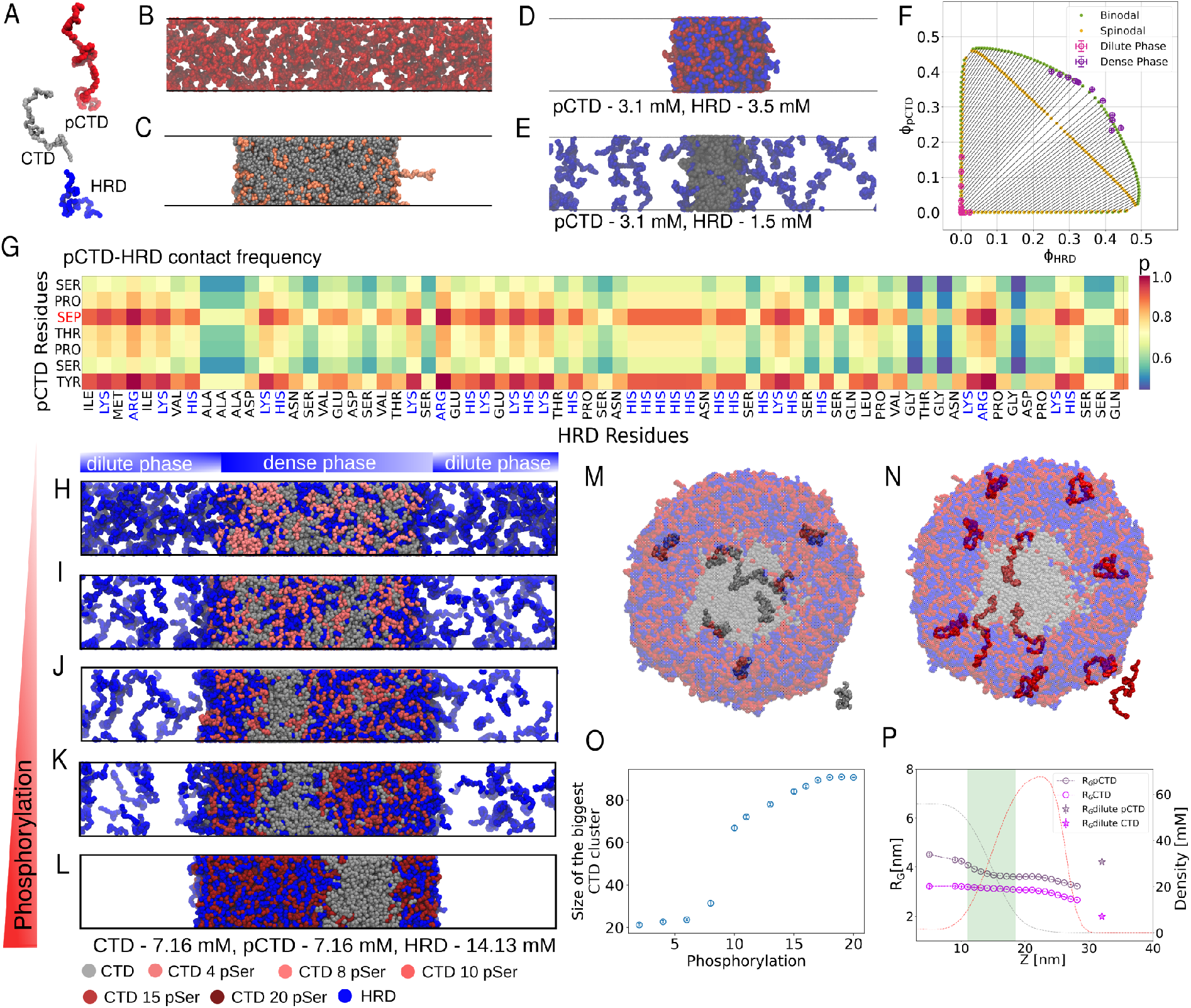
Co-phase separation HRD with CTD and pCTD in slab simulation. A) Single chains of CTD, pCTD, and HRD. B) Fully phosphorylated CTD disrupt the phase behaviour. C) CTD condensate with 50 partially phosphorylated pCTD chains with 10 pSer_5_ phase separating together.2 D) Simulation snapshot showing pCTD and HRD forming condensate. E) Simulation snapshot of CTD with HRD. While CTD forms a condensate, HRD exhibits no favourable interaction with CTD. F) Biphasic diagram of the pCTD-HRD system. Positive tie lines indicate associative phase separation. The volume fractions from molecular dynamic simulations are in magenta and purple. Binodal and spinodal from mean-field fit are shown in green and yellow dots. G) Contact frequency analysis between pCTD and HRD showing the important electrostatic interactions between positive (blue) and negative (red) charged residues. H-L) Mixing and de-mixing of the dense phase of 100 CTD, 100 pCTD, and 250 HRD chains, with increasing the degree of pCTD phosphorylation. H) 4pSer I) 8pSer J) 10pSer K) 15pSer L) 20 pSer of pCTD are phosphorylated. M) Conformation of CTD and pCTD within the droplet. Snapshot capturing CTD chain conformations, where the *R*_*G*_ of CTD chains reduces slightly in the pCTD-HRD phase. In the dilute phase, the chains become more compact. N) Snapshot showing pCTD chain conformations. The chains are more elongated in the CTD phase and compact in the pCTD-HRD phase. The chains are elongated in the dilute phase. O) The size of the biggest CTD cluster as the phosphorylation of pCTD varies in the simulation. P) Comparison of the Variation in *R*_*g*_ (radius of gyration) between CTD and pCTD Conformations, Analysed from the Center of the Condensate.

To explore the impact of different patterns of distributing an identical number of phosphorylated Ser_5_ residues on CTD sequences, we conducted simulations with partially phosphorylated pCTD chains, each carrying different distribution of pSer, in the presence of a CTD condensate and compared the variation in dilute concentration of pCTD (Fig S7 C). We considered five distinct distributions: maximally distributed in every second heptad repeat, in adjacent heptad repeats, in the middle, and in N and C-termini, respectively. For pCTD chains with 10 phosphorylation values (below the threshold), rarely any pCTD chains were found outside the CTD condensate, regardless of their distribution. With 15 pSer (above the threshold), the dilute concentration remained nearly the same value for different distributions. In conclusion, our simulations show a phosphorylation dependent demixing of unphosphorylated and phosphorylated CTD. The degree of demixing is thereby dependent on the number of phosphorylation sites, but not their distribution along the CTD heptad repeats.

Thus, our simulations can explain why in vitro CTD condensates are dissolved over time in the presence of the kinase CDK7 and ATP^13^. However, this also poses a clear contradiction with all observations made in cells^14,23^, where transcribing RNA Polymerase CTD localizes into condensates even when it is highly phosphorylated^15^. A likely explanation for this discrepancy is that in cells the phosphorylation of CTD can create binding sites for multiple other interacting proteins^14^. We therefore wondered if we can simulate the impact of additional factors in our model. As a first indicator, we consider the interactions of CTD and pCTD with FUS condensates. In our simulations the unphosphorylated CTD binds more strongly to FUS low complexity domain (LCD) condensates than pCTD (Fig S8), which is in line with experiments that demonstrated that CTD phosphorylation abrogates the binding of CTD to FUS gels and condensates. Interestingly while FUS and CTD both localize to a single dense phase, they are not homogenously mixed in the condensate. We further observed that FUS results in a minor enrichment of pCTD in the dense phase. Together this shows that we can indeed expand our simulation to encompass CTD interaction partners.

### Co-phase separation pCTD and HRD by complex co-acervation

FUS mainly interacted with the non-phosphorylated form of CTD. Inside the cell, numerous interaction partners of CTD might shield the effect of CTD phosphorylation. One protein with a well described role in transcriptional elongation is Cycline T1. As part of the positive transcription elongation factor b (P-TEFb) Cycline T1 stabilises elongating RNA Polymerase II and it is essential for pCTD condensation in vitro and potentially also in vivo. In agreement with experiments, simulations show that pCTD co-phase separates with the histidine rich domain (HRD), which is a disordered domain of P-TEFb Cyclin T1^14^.

HRD, which is positively charged on account of its histidine side chains, on its own does not interact strongly with CTD (Fig 3E). CTD chains in our simulation are uncharged since we are simulating an idealized version of the CTD with no deviation from the canonical heptad Y_1_S_2_P_3_T_4_S_5_P_6_S_7_ repeat motifs. In the pCTD and HRD condensates (Fig 3 D), pSer residues of pCTD interact strongly with positively charged residues Arg, Lys, and His in HRD. pSer forms few contacts with Ala, Ser, Thr, and Gly residues of HRD (Fig 3 G). Tyr in pCTD also engages in strong contacts with Lys and Arg in HRD and to a somewhat lesser degree with many other residues, e.g., Ile, Met, and Val, but interacts only weakly with Ser and Gly. The contact analysis clearly shows that context matters a lot^70^, a negatively charged Asp next to a positively charged Lys residue is more frequently in contact with pSer than uncharged Ala residues in a stretch of Ala residues.

To understand the interaction between pCTD and HRD, we conducted simulations to construct a phase diagram, which shows that the pCTD and HRD form complex coacervates through electrostatic interactions. Throughout these simulations, we kept the concentration of HRD constant at 3.9 mM while systematically increasing concentrations of pCTD (Fig S9). Considering that pCTD and HRD are oppositely charged we expect them to interact electrostatically and form a complex coacervate. This type of phase separation, also known as cross-interaction-driven phase separation (CIP), happens when two species are mutually attracted to each other and cannot phase separate independently^71^. CIP typically gives rise to a looped phase diagram, since the binodal lines do not intersect with the composition axes. The relative concentrations of the proteins are critical for determining the extent of phase separation and the location of the coexistence region. Notably, a rise in the concentration of either protein can initiate phase separation, leading to the formation of condensates. This occurrence is depicted in Fig S8 E, where it is shown that, at an approximate concentration of around 5 mM of pCTD, all the HRDs in the dilute phase undergo phase separation. Further increasing the pCTD concentration has no effect on the dense phase, rather, the dilute concentration of pCTD increases. The saturation concentration of HRD gradually decreases with increasing pCTD concentration, as HRDs initially present in the dilute phase transition into the dense phase by associating with added pCTD chains. This observation becomes even clearer when examining the ternary phase diagram depicted in Fig 3F. Here, the pink and purple dots represent the simulated coexisting (binodal) dilute and dense phases respectively. Phase separation is associative, evidenced by the positive tilt in the (imaginary) lines that connect the coexisting phases in composition space (tie-lines). This is further supported by fitting the simulated binodal compositions in Fig 3F using Flory-Huggins theory (Eq. 4) (SI Text), which is shown by green dots. The fit also strengthens our interpretation that self interactions of hyper-phosphorylated CTD and HRD are negligible due to electrostatic repulsion. It is consistent with the observation that the dilute phase is devoid of polymers of one type in the simulations, but enriched by polymers of the other type if their average volume fraction increases, which is also expressed by the tie-lines which develop an off-set at either axis as soon as the mean composition ratio starts deviating from unity (Fig 3F, grey).

We conclude from the simulations and from fitting the Flory-Huggins model to the simulation data that, unlike CTD condensates where Tyr-Tyr and Tyr-Pro contacts drive phase separation, pCTD, and HRD form co-acervates by electrostatic interactions, which suggests that on account of differences in molecular driving forces CTD and pCTD-HRD phases could in principle constitute distinct environments in the nucleus. The excellent fit with the simulated data, despite the fact that Flory-Huggins theory does not include a description for (long range) electrostatic interactions, supports considerable screening by the implicit background salt in the simulations, making the attractive forces between the polymers effectively short-ranged. Furthermore, owing to the predominance of a single type of interaction, a more complex mean-field model for fitting the data^72^ would be redundant. The overall attraction between the polymers is captured by a negative polymer-polymer interaction parameter: *χ*_*AB*_ = −2.36. The polymer-solvent interaction parameters (see Eq. 4), as obtained from the fitting, are *χ*_*AS*_ = 0.55 and *χ*_*BS*_ = 0.36, indicating relatively poor stabilisation (near theta conditions) of, in particular, the HRD.

### Phosphorylation can act as a switch and trigger the formation of a second dense phase in the co-phase separation of CTD, pCTD, and HRD

In separate simulations, increased phosphorylation disrupts the phase separation of pCTD. However, the binding of HRD to pCTD facilitates its phase separation. Yet, within cells, CTD, pCTD, and HRD coexist concurrently, raising the question if they form a single mixed phase or segregate into two distinct dense phases.

Thus we have simulated fixed numbers of CTD, pCTD, and HRD and increased the degree of pCTD phosphorylation in a series of simulations. Partially phosphorylated pCTD containing 4 pSer_5_ co-phase separates with CTD, as it is apparent from the density profile (Fig 3H, S10). The density profile illustrates that both pCTD, CTD, and HRD chains are present within the mixed condensate, while a significant portion of HRD is in the dilute phase. However, when the degree of phosphorylation of pCTD increases and 8 Ser_5_ are phosphorylated (Fig 3I) the amount of HRD in the dilute phase decreases as it is now recruited to the dense phase by the pCTDs. There is a tendency for pCTD and HRD to co-localize as apparent in the density profiles. With 10 pSer_5_ phosphorylated, the HRD concentration in the dilute phase is markedly reduced further and the co-localization of pCTD and HRD is enhanced (Fig 3J). The co-localization of pCTD-HRD is observed to favour the periphery of the condensate border (Fig 3J-L). This preference for condensate periphery localisation could be attributed to the presence of negative charges on pCTD and positive charges on HRD, potentially facilitating interactions with the solvent compared to the uncharged CTD chains. With further increasing phosphorylation of pCTD, there is an enhanced tendency for both pCTD and HRD to localise towards the boundaries of the CTD-pCTD-HRD condensate. Unphosphorylated CTD is clearly at the centre of the condensate. Adding more phosphates to pCTD results in a core-shell morphology, with the CTD density mostly at the centre of the condensate, with steep decreases in CTD density towards the boundaries. De-mixing is not complete, since some CTD chains co-localise with pCTD-HRD as is apparent from both the visualisation of the simulation and the density profile. Phosphorylating all Ser_5_ residues in the 140mer pCTD chains (Fig 3L), further enhances the de-mixing.

The size of the largest CTD cluster in simulation shows a sharp transition when about half of the Ser_5_ of pCTD chains are phosphorylated, indicating the onset of the separation process into two distinct phases: CTD and pCTD-HRD (Fig 3O). The microphase separated domains of the homopolymer CTD phase and the heteropolymer pCTD-HRD phase begin to evolve as the degree of phosphorylation increases. This change becomes more pronounced with increasing phosphorylation, leading to an expansion in the size of these domains. The excess of CTD in the vicinity of CTD and pCTD (*n*^*exc*^) also shows a similar transition as phosphorylation increases (Fig S10 F). The parameter, *n*^*exc*^, reveals how the number of nearby pCTD and CTD chains within a specific distance from a CTD chain changes with increasing phosphorylation of pCTD. It shows changes in the local environment, providing insights into the frequency and strength of interactions between CTD and pCTD chains as the phosphorylation of pCTD varies. At lower phosphorylation levels, a CTD chain is surrounded by both pCTD and CTD, and the numbers of CTD and pCTD are consistent. However, a noticeable deviation becomes apparent as the degree of phosphorylation surpasses 10 pSer_5_.

Additionally, the overall structure and shape of pCTD chains are affected by phosphorylation (Fig S11). In dilute solution, phosphorylation expands the protein in line with experiments^73^ as the chains become less globular and more rod-like due to electrostatic repulsion. Similarly, in the dense phase, pCTD also adopts a more rod-like conformation. The radius of gyration (*R*_*G*_) linearly increases with phosphorylation below a threshold of 10 pSer_5_. Once this threshold is crossed, *R*_*G*_ stabilises and the anisotropy of pCTD chains decreases as interactions with HRD compensate for electrostatic repulsion. This shift once again highlights the impact of phosphorylation in modulating specific interactions between the protein pairs in forming two distinct dense phases.

The conformations of CTD and pCTD are modulated by the different phases and interfaces within the condensate (Fig 3 M, N) as the CTD and pCTD-HRD phases indeed have different properties. These differences could underpin distinct regulatory processes mediated by the two phases within a biphasic condensate. In a “good” solvent, which is compatible with the chain, the chain is extended, whether the solvent is a dilute solution or a dense phase. By contrast, a chain tends to collapse in a “poor” solvent which is not compatible with a chain. CTD is more expanded in the condensate than in a dilute aqueous solution, as tracked by the radius of gyration (*R*_*G*_) (Fig 3P, S12 A, and SI Text). The aqueous phase is a poor solvent for CTD chains since the ideal YSPTSPS heptads are devoid of charged groups and feature aromatic and non-polar residues. The conformation of pCTD is modulated by the CTD and pCTD-HRD phases in a different way. Hyperphosphorylated pCTD is more extended in the dilute aqueous phase than in the condensate (Fig S12 B).

### Morphology of CTD and pCTD condensates in simulations, in vitro, and *C. elegans* embryos

The simulation reveals a core-shell arrangement and by calculating interfacial tensions through our molecular dynamics simulations^26,74^, we examined the potential morphologies these condensates could take on and explained the reasons why such a core-shell structure is possible. When two immiscible droplets of CTD and pCTD-HRD are brought into contact, four distinct morphological configurations become possible^6,75^. We visually represent these four distinct morphologies using a stability diagram generated from the contact angle equation, as depicted in Fig 4D. The contact angle equation is given by:

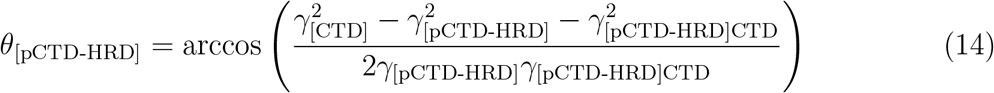

A schematic shows partially engulfing CTD and pCTD-HRD phases, illustrating the contact angles between these phases in Fig 4D. The stability diagram describes the four different morphologies attainable by adjusting relative interfacial tensions, and these variations can be characterised through the corresponding contact angles. In Region I, the CTD phase is entirely enveloped by the pCTD-HRD phase, with the condition *γ*_[CTD]_ > *γ*_[pCTD-HRD]_ + *γ*_[pCTD-HRD]CTD_. In Region II, the CTD and pCTD-HRD phases are partially engulfed, sharing an interface while also being exposed to the dilute phase. The transition from complete engulfment to non-engulfing phases is gradual as *θ*_[pCTD-HRD]_ increases. Here, the three interfacial tensions are comparable: *γ*_[CTD]_ ∼ *γ*_[pCTD-HRD]_ ∼ *γ*_[pCTD-HRD]CTD_. In Region III, the significant interfacial tension between the CTD and pCTD-HRD droplets leads to a non-engulfing morphology, defined by *γ*_[pCTD-HRD]CTD_ > *γ*_[CTD]_ + *γ*_[pCTD-HRD]_. In Region IV, the CTD phase engulfs the pCTD-HRD phase. The transition from Region 4 to Region I is marked by a black arrow, signifying an increase in *θ*_[CTD]_.

**Figure 4.**
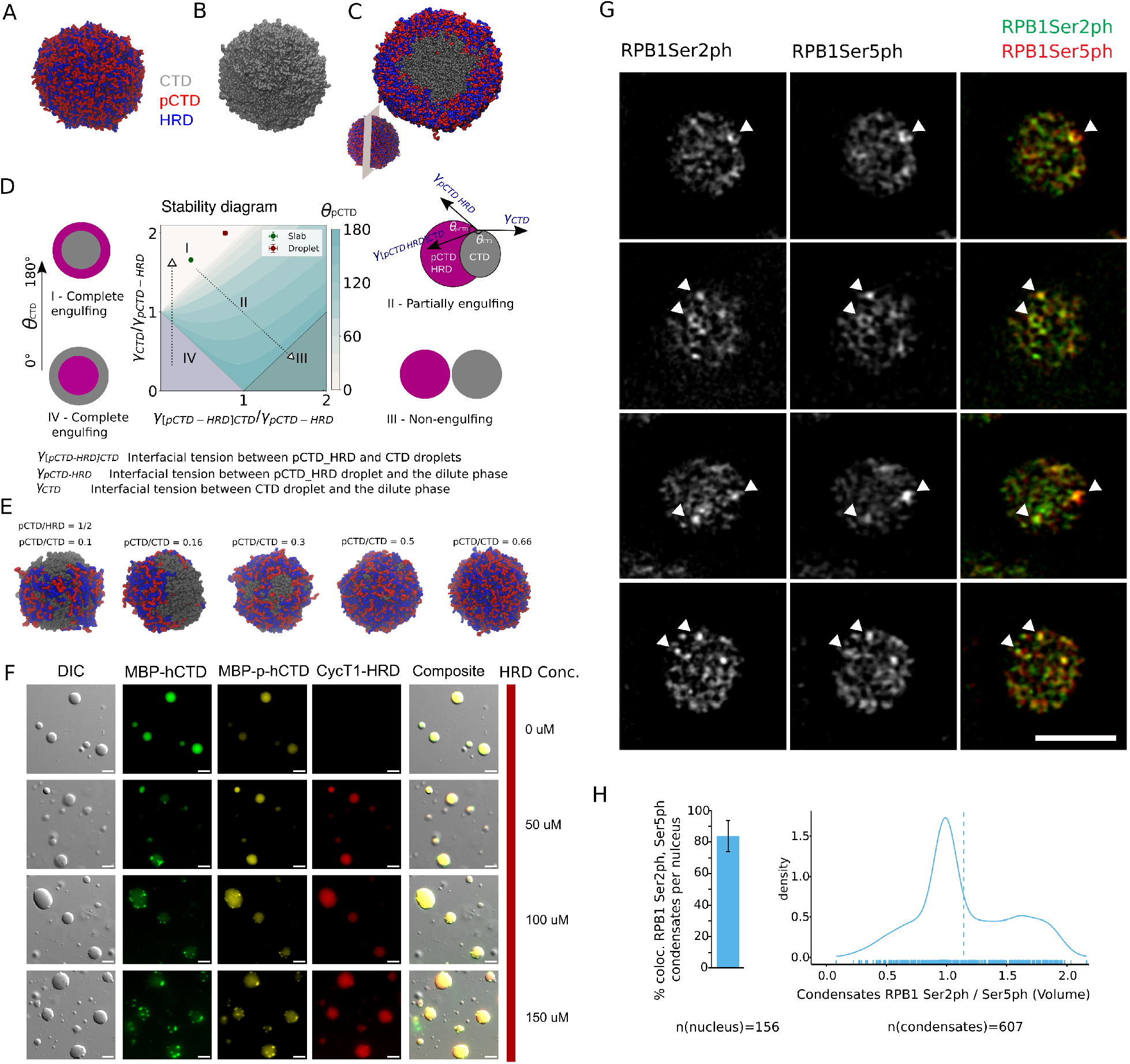
Morphology of the multiphasic condensate A) Formation of a droplet composed of 900 pCTD and 1800 HRD chains. B) Droplet comprising 600 CTD chains. C) Multiphasic droplet with a core-shell morphology, constituted by 900 pCTD, 1800 HRD, and 600 CTD chains. D) Stability diagram illustrating four distinct morphologies arising from the interaction between CTD and pCTD-HRD droplets. Interfacial tension analysis from slab and droplet simulations indicates the core-shell morphology. Sketch illustrating a partially engulfing CTD and pCTD-HRD phases indicating the contact angles. E) CTD condensate partially enveloped by pCTD and HRD condensate. Sequential images depict the progression from partially engulfed to fully engulfed morphology as the ratio of pCTD/CTD chains increases. F) Droplet compartments induced by HRD. Differential interference contrast (DIC) shows MBP-hCTD (AF488; green), MBP-p-hCTD_partial_ (AF594; yellow) and HRD (AF647; red) forming different protein condensates upon the addition of HRD. Composite pictures show droplets formed by different components in function of HRD’s concentration. G) Super-resolution microscopy image of *C. elegans* embryos stained with antibodies targeting phospho-Serine 5 (pSer5, indicating transcriptional initiation) and phospho-Ser2 (pSer2, indicating transcriptional elongation) phosphorylated RPB1. RPB1 labelled with either PTM is localized in prominent subnuclear foci. H) The segmentation and quantification of these foci demonstrate that both types of foci colocalize, exhibiting only partial overlap.

Full engulfment of CTD but pCTD-HRD is the most favourable morphology, but small changes in the system would enable partial engulfment. We used capillary wave theory (CWT)^74^ to compute the interfacial tensions of CTD and pCTD-HRD with slab geometry with each Ser_5_ in the heptad repeats of pCTD phosphorylated. Three sets of slab simulations were performed to determine interfacial tension at different interfaces. The first two simulations involved CTD and pCTD-HRD being simulated independently, with each forming a condensate, from which the interfacial tension at the interface with the dilute phase can be calculated. In the third simulation, CTD and pCTD-HRD were simulated together, forming a multiphasic condensate, and we computed the interfacial tension at the interface between the CTD and pCTD-HRD condensate (SI Text and Fig S13). Evaluating the ratio of interfacial tension for different interfaces, specifically *γ*_[CTD]_*/γ*_[pCTD-HRD]_ and *γ*_[pCTD-HRD]CTD_*/γ*_[pCTD-HRD]_, we confirmed that the observed core-shell morphologies are consistent with the theory of liquid interfaces (depicted by the green dot in Fig 4D). This supports our conclusion that the core shell morphology in our simulations is stable state and our observation is not an artefact of insufficiently long simulations. To rule out effects from the box geometry, we repeated the simulations in large cubic boxes 150 nm×150 nm×150 nm with up to 3300 proteins chains, where the proteins form spherical droplets (Fig 4A-C, Movie S2). The ratios of interfacial tension between different interfaces, *γ*_[CTD]_*/γ*_[pCTD-HRD]_ and *γ*_[pCTD-HRD]CTD_*/γ*_[pCTD-HRD]_, were computed from the shape fluctuations method^26^ (indicated by the red dot in Fig 4D). These results further confirm that the core-shell morphology is the most favourable arrangement for the CTD, pCTD-HRD system in our simulations. The comparison of interfacial tension at different interfaces using various methods is illustrated in Fig S14, demonstrating consistent values across the different approaches. We note that the computed ratios of surface tensions are closer to the border between full and partial engulfment. We can also observe partial engulfment of CTD in simulations where there are not enough pCTD and HRD chains to fully cover the CTD surface (Fig 4E, Movie S3). Simulations clearly show that the condensate of pCTD-HRD either fully or partially surrounds the CTD condensate (Fig 4C, E). At lower concentrations of pCTD and HRD, they undergo coacervation and form a small patch on the surface of the CTD condensate rather than spreading evenly across the surface, resulting in a partially engulfed structure. With an increase in the concentration of pCTD and HRD, the degree of engulfment also increases, indicating that higher concentrations of pCTD and HRD lead to a decrease in interfacial tension between the pCTD-HRD complex and CTD^76^.

We asked whether such composition-dependent partial demixing of CTD, pCTD, and HRD can also be observed in vitro. Upon addition of CDK9/Cyclin K to human CTD (h-CTD), we obtain partially phosphorylated CTD (p-hCTD_partial_), with up to nine chains featuring phosphates. (Fig 4F, S15)

HRD on its own does not phase separate even at high concentrations. Confirming predictions in simulations, the mixture of pCTD_partial_ (MBP-tagged) and HRD recovers the phase transition in a concentration-dependent manner (Fig 3). At low and mid molar ratios (1:1:1-6; hCTD:p-hCTD_partial_:HRD) hCTD tends to recruit both proteins but some puncta are observed with a high content of hCTD plus a gradual decrease of green in the big droplets consistent with an enrichment of pCTD_partial_ and HRD. This is in line with what we observe in simulations (Fig 3 H,I). The morphology in vitro does not show engulfment of CTD. This is due to the lower phosphorylation level of pCTD, which allows it to interact with both phosphorylated and unphosphorylated CTDs. This will lower *γ*_[pCTD−HRD]CTD_ and thus favour partial engulfment, where CTD might be at the surface of a p-hCTD_partial_-HRD-CTD droplet. The last ratios used (1:1:8-10), with higher levels of HRD than in simulations, show a complete exclusion of hCTD from some of the droplets observed by DIC in agreement with a partial exclusion of p-hCTD. Surprisingly, HRD remains almost uniformly distributed. Here, the partition of the lowly-phosphorylated p-hCTD_partial_ is assumed to play a major role where the condensates excluding hCTD are potentially enriched in p-hCTD_partial_/HRD mixtures, as expected for gene-body or elongation condensates, while the droplets enriched in hCTD would support the initiation stage of transcription.

The simulations thus suggest that RNA Polymerase II condensates in cells might be segregated into partially, or fully engulfed condensates, which we test in vivo by super-resolution microscopy of *C. elegans* embryos. We stained the *C. elegans* embryos with antibodies against phospho-Serine 5 (pSer_5_, as a marker for transcriptional initiation) and phospho-Ser2 (pSer_2_, as a marker for transcriptional elongation) phosphorylated RPB1. RPB1 marked with either PTM can be found in large subnuclear foci (Fig 4G). Segmentation and quantification of these foci revealed that both foci types colocalize, but only partially overlap (Fig 4H). This is consistent with the partial or full engulfment of two distinct condensates, as seen in the simulations. Indeed a fraction of the pSer_2_ and pSer_5_ RPB1 foci formed hollow spheres, or half-moon like structures (Fig 4G). Given that total RPB1::GFP formed solid round foci (Fig 2E and Ref^77^), we conclude that these hollow spheres are part of a bigger RNA Pol II condensate. This highlights partial and full engulfment of CTD by pCTD in remarkable agreement with our simulations and hints at the prevalence of patterns of RNA polymerase II in different phosphorylation states.

Consistent with our simulations CTD condensates are stable *in vivo*, even when Cyclin T1 is knocked down by RNA interference (RNAi) and Ser_2_ phosphorylation is consequently abolished^78^ (Fig S16). This shows that CTD condensates can form in the absence of Ser_2_ phosphorylation as in our simulations. The condensates are generally more round than in the presence of Cyclin T1. A more regular round appearance is consistent with the more spherical shapes of CTD condensates in contrast to more irregularly shaped condensates of CTD, pCTD, and HRD.

### Recruitment of FUS and partially phosphorylated CTD to multiphasic condensates

Having established that the two phases of condensates of CTD, pCTD, and HRD are distinct chemical environments, we investigated whether they could indeed differentially interact with other molecules, which would enable each phase to underpin different stages of transcription. We observed contrasting interaction patterns between CTD and pCTD-HRD with phase-separated LCD FUS. CTD interacts with phase-separated FUS as apparent from snapshots of simulation systems of CTD, pCTD-HRD, and FUS (Fig S17). Regions of the simulation system enriched in pCTD-HRD are relatively depleted in FUS. This preference for the CTD phase over the pCTD-HRD phase is consistent with the proposed roles of CTD chains and CTD condensates in supporting transcription initiation by interacting with transcription factors to form promoter condensates and the experimentally established preference for FUS LCD to interact with CTD over pCTD^79,80^. We checked the partitioning of partially phosphorylated pCTD to multiphasic condensates and found that partially phosphorylated CTD can interact with both the CTD and pCTD-HRD phases (Fig S18). The partially phosphorylated CTD is relatively depleted at the peaks of the pCTD and HRD concentration profiles and shows how a partially phosphorylated CTD can reside in the condensate phase associated with initiation and the RNA polymerase II would only switch to the other phase upon further phosphorylation.

## DISCUSSION

We show how phosphorylation of CTD can give rise to distinct compartments which could underpin transcription regulation. Previously, it was shown that perturbation of transcription condensates impairs transcription^81^ and in vivo IDRs which can form liquid condensates have been shown to be instrumental for transcription^61^. Experiments have suggested that transcription initiation and elongation may be regulated in time and space by phase-separated condensates^15,77^. How this may be achieved by proteins enriched in seemingly structured and feature-less IDRs^82–84^, has remained elusive. With molecular dynamics simulations, we show how the interactions of CTD and pCTD with additional proteins such as HRD of Cyclin T1 which can shield the negative charges introduced by hyper-phosphorylation can give rise to distinct phases, one defined by hydrophobic interactions and the other phase by electrostatically driven cross-interaction driven phase separation. For the formation of entirely distinct phases, pCTD phosphorylation needs to cross a threshold upon which the interactions of CTD and pCTD change drastically. In phase-separation experiments with purified proteins, where pCTD is partially phosphorylated we observe partial demixing of CTD from p-hCTD_partial_ and HRD (Fig 4), which is in line with our simulations (Fig 3 I-J). This rationalises how highly phosphorylated pCTD of elongating RNA Polymerase II in zebrafish is excluded from CTD clusters and condensates in zebrafish embryos^23^. CTD is fully and partially engulfed by pCTD-HRD in our simulations. For a given composition of the simulation system, we determine its most favourable morphology by computing interfacial tensions.

Our approach is completely general and can be extended to any kind of mixture of proteins. Thus it becomes possible to predict the morphologies of multi-component mixtures from molecular simulations. The coarse-grained model we employed to simulate large condensates captures the contact statistics of CTD as reported by NMR and atomistic molecular dynamics, is in line with interactions propensities for CTD chains from atomistic molecular dynamics and also reproduces experimentally determined effects of truncations and sequence changes.

In accordance with our simulations, super-resolution microscopy of *C. elegans* embryos reveal full and partial engulfment of CTD by pCTD. RNAi knockdown of Cyclin T1 preserves CTD condensates, which is in line with the stability of CTD condensates in the absence of Ser phosphorylation suggested by our simulations (Fig S16). While the two condensates are distinct and modulate the global conformations of proteins in different ways, the two dense phases are in contact (Fig 4). The two phases also differentially recruit FUS as expected, with FUS binding preferentially to the CTD rather than pCTD-HRD. Thus our findings provide a molecular basis for the observations by Wei et al., who report that in cells Ser_5_ phosphorylated CTD concentrically organises around a TAF-15 promoter condensate^77^. While the study by Wei et al., did not probe the makeup of the pCTD shell, whether pCTD accumulated on the exterior of the TAF-15 condensate on its own or in a dense phase, this core-shell-like accumulation of pCTD around a promoter condensate is reminiscent of the morphologies we have resolved in simulations and in cells and suggest that a promoter condensate may be engulfed by a gene-body condensate.

Our study highlights an additional and unexpected layer of specificity which CTD condensates may impart^19^. Experiments showed that CTD forms condensate in a temperature dependent manner in *C. elegans* embryos. Many protein condensates dissolve at high temperatures. Yet transcription rates do not decrease with moderate temperature increases. On the contrary, transcription is typically enhanced. This apparent conundrum is resolved by our *in vivo* experiments, simulations, which highlight that the LCST behavior of CTD would be consistent with the involvement of condensates in transcription and an increase of transcription at elevated temperatures. The LCST behaviour of CTD, which has previously been characterised in biophysical experiments^22^, correlates with moderate shifts in gene transcription programs with temperature as shown by our RNA sequencing data. The LCST behaviour is largely uncoupled from a classical heat shock response, which results in dramatic changes in the pool of RNA transcripts going from 30 °C to 37 °C. At temperatures below 37 °C changes in the phase separation propensity of CTD in *C. elegans* correlate with a fine-tuning of gene transcription as temperature changes, with most changes seen for transcripts coding for proteins in metabolic pathways, which may be an important facet of homeostasis in non-stress conditions. This LCST behaviour of CTD contrasts with the temperature behaviour of other proteins in the nucleus, such as the heterochromatin proteins MET-2^57^ and HP1*α*^69^, which dissolves at higher temperatures. These differences further highlight that CTD condensates that support transcription are physio-chemically distinct from heterochromatin condensates, which is likely essential for transcription regulation.

## ACKNOWLEDGEMENTS

A.C.S is supported by M^3^ODEL. A.C.S, F.S. and L.S.S. thank M^3^ODEL for support. L.S.S. acknowledges support by ReALity (Resilience, Adaptation and Longevity) and Forschungsinitiative des Landes Rheinland-Pfalz. This project was funded by SFB 1551 Project No. 464588647 of the DFG (Deutsche Forschungsgemeinschaft). Funding from the German Science Foundation is acknowledged: F.S. and L.S.S. are members of the RTG 2516 (Grant No. 405552959). Further, we gratefully acknowledge the advisory services offered and the computing time granted on the supercomputers Mogon II at Johannes Gutenberg University Mainz, which is a member of the AHRP (Alliance for High Performance Computing in Rhineland Palatinate) and the Gauss Alliance e.V. We thank Dr. J. Mittal, Dr. P. Virnau, Dr. M. Andrade, Dr. G. Hummer, and Dr. R. V. Pappu for inspiring discussions and Dr. G. Krawezik for help with the atomistic simulations on Folding@Home. STELLARIS 8 Falcon confocal was funded by Deutsche Forschungsgemeinschaft (DFG, German Research Foundation) project number 497669232. All imaging and RNA sequencing was supported by the IMB Microscopy Core Facility and IMB Genomics Core Facility.

