## Supporting Information for "Promoter and Gene-Body RNA-Polymerase II co-exist in partial demixed condensates"

[5)] *Max Planck Insitute for Polymer Research, Ackermannweg 10, 55128 Mainz, Germany*

[6)] *Institute of Molecular Biology (IMB), 55128 Mainz, Germany*

[7)] *Flatiron Institute, New York, USA*

[8)] *Department of NMR-based Structural Biology, Max Planck Institute for Multidisciplinary Sciences, Am Faßberg 11, D-37077 Göttingen, Germany.*

[9)] *KOMET1, Institute of Physics, Johannes Gutenberg University Mainz, Staudingerweg 7, 55128 Mainz, Germany*

[a]Electronic mail:
[b]Electronic mail:
[c]Electronic mail:
[d]Electronic mail:

### Comparison of atomistic and coarse-grained simulations

Dissociation constants ($K_d$) are often used to measure binding interactions, which here we estimated from a kinetic-rate model[1,2]. The dissociation constant of two proteins can be directly estimated from the binding probability, $P_b$, which indicates the proportion of frames where the chains are in the bound state. We can use the following equation to estimate the dissociation constant[3]:

$$K_d = \frac{(1 - P_b)^2}{N_A V P_b}, \tag{1}$$

where, $N_A$ is Avogadro's number and $V$ is the box volume. $P_b$ is determined from the simulation involving two peptides.

By analyzing the simulations in terms of a kinetic-rate model[1,2] (time-continuous Markov-state model), we can derive more accurate estimates for the relative populations of different states in the simulations, in this case $P_b$ and $1 - P_b$. In a kinetic model, the time spent waiting in the different states and the number of transitions from one state to another are key inputs and can thus enable accurate estimates of relative populations even when analysing relatively short trajectories[4,5]. We use a dual cutoff scheme for the time series data generated from the simulations to track the transitions between the bound and unbound states of the protein chains and estimate the probability of observing the bound state. This dual cut-off scheme is also referred to as transition-based state assignment[6] (Fig S1A,B). The "Dynamic Histogram Analysis Method extended to detailed balance" (DHAMed) package (https://github.com/bio-phys/PyDHAMed) is used to calculate the probability of the bound and unbound states $P_b$ and $1 - P_b$. Since detected many transitions between bound and unbound conformations in 400 short all-atomic simulations on Folding@Home it is reasonable to estimate $K_d$ values from these simulations and use these to compare the all-atom and coarse-grained molecular dynamics simulations. The magnitude of the $K_d$ is similar to what has been reported by Tesei et al.[7]. In the case of two CTD chains all-atom simulations, we obtained $K_d = 2.76 \pm 0.24$mmol, whereas in the pCTD-CTD simulations, the $K_d = 1.87 \pm 0.20$ mmol.

Considering the frequent transitions between bound and unbound states and thus overall reasonable sampling of our simulations, we also computed the radial distribution function from 39.8 µs atomistic trajectories for two 21 residue CTD interacting, where water and ions are represented explicitly and also from coarse-grained simulations of such a fragment. The two fragments overall interact in a similar way as captured by evaluating the second virial coefficients. Note that the numbers we report here are $B_{22} = -0.81\mu$l mol g$^{-2}$ and $B_{22} = -0.57\mu$l mol g$^{-2}$ for CG and all-atom respectively, are similar to what Tesei et al. have reported for the interactions of disordered FUS protein low complexity domain (LCD) chains[7].

These calculations are based on the Mayer integral for the second virial coefficient:

$$B_{22} = \int_0^\infty 2\pi r^2 [\exp(-U(r)/k_B T) - 1]\, dr, \tag{2}$$

where $U(r)$ is the total interaction potential between the two fragments. At high dilution, one can approximate $\exp(-U(r)/k_B T) \approx g(r)$ and thus calculate the second virial coefficient from $g(r)$:

$$B_{22} = \int_0^\infty 2\pi r^2 [g(r) - 1]\, dr, \tag{3}$$

which are a measure of how much molecules like to interact with each other.

### Contact analysis of atomistic and coarse-grained simulations

Contact maps were computed with the contact-map-explorer Python library (`https://github.com/dwhswenson/contact_map`). For coarse-grained and atomistic simulations, a cut off of $2^{1/6}(\sigma_i + \sigma_j)/2$, where $\sigma$ is the diameter of the residues, was employed to define whether a pair of residues are in contact in a given simulation frame.

### Mean-field theory analysis of phase behaviour of pCTD and HRD

The phase behavior of binary solutions of biopolymers can, perhaps conveniently, often be captured by mean-field theory, irrespective of knowing the exact nature of the underlying interaction pattern[8]. In what follows we show that the mixture of HRD and pCTD in buffer forms no exception with this respect. A theory-based interpretation of experimental or simulated data is useful, as it demonstrates in what way the complexity may be reduced to an effective parametrization of a simplified model, while at the same time identifying the relative importance of the different interactions involved. Here, we fit the simulated binodal compositions in Fig 4J using Flory-Huggins theory, which gives the dimensionless Helmholtz free energy density for the ternary mixture as in Eq. 4[9].

We obtained the values for the interaction parameters from fitting the binodal curve (green) of a phase diagram calculated using Eq. 3 to the simulated coexisting compositions. An excellent fit with the simulated data is obtained, despite the fact that Flory-Huggins theory does not include a description for (long range) electrostatic interactions. The latter might mean that the background salt, implicitly assumed in the coarse-grained simulations, makes the attractive forces between the polymers relatively short range. The fit values for the interaction parameters are: $\chi_{AB} = -2.36$, $\chi_{AS} = 0.55$ and $\chi_{BS} = 0.36$. The highly negative polymer-polymer interaction parameter (effectively) captures the energy balance associated with the electrostatic attraction between HRD and pCTD, in combination with the repulsion between like species. The polymer-solvent interaction parameters are both below their critical values, $\chi_c = \frac{1}{2}(\frac{1}{\sqrt{N_{i=A,B}}} + 1)^2$ (= 0.63 and 0.59 for HRD and pCTD, respectively), a prerequisite for a looped miscibility gap. We note that $\chi_{AS}$ lies closer to its critical value than $\chi_{BS}$, suggesting the pCTD to be somewhat better accommodated by the salt solution than HRD. Finally, we note that we can obtain a fit of similar quality with a different set of fit parameters, where $\chi_{AB}$ is slightly less negative, and this is compensated by a modest increase of $\chi_{AS}$ and $\chi_{BS}$. However, this set of parameters would not reproduce the full miscibility loop, therefore we discard it.

The fact that a fit to the simulation data can be obtained is also consistent with the notion that the simulation runs are long enough. For short simulation runs, we would expect that concentrations of the two phases to be strongly affected by noise and thus an overall poor fit would be expected. The good agreement then enables us to proceed to study the phase separation of pCTD at different levels of phosphorylation in the presence of HRD and CTD.

#### Characterisation of pCTD structure in pCTD-HRD condensates

We can use the simulations to better understand how phosphorylation modulates the structure and shape of pCTD chains in dilute and dense phases. For single chains of pCTD in the dilute phase, we observed an increase in $R_G$ and anisotropy with increasing phosphorylation (Fig S11 A, B and inset), indicating a conformational change from a globular-like to an elongated structure. Small-angle X-ray scattering experiments have previously established that phosphorylation of dilute solution of CTD from *Drosophilia* leads to an increase in $R_G$ values[10]. Swelling of CTD upon phosphorylation has also been observed in atomistic molecular dynamics simulations[11]. The addition of phosphates to CTD initially results in increased asymmetry among the chains, due to repulsion between different parts of the chains. However, upon analyzing the radius of gyration ($R_G$) values of pCTD chains in the dense phase, we observe only slight deviations compared to those in a sparser environment, as depicted in (Fig S11B). At lower phosphorylation levels, pCTD exhibited comparable affinities for both CTD and HRD. A slight increase in $R_G$ and the anisotropy of the chains is observed as the phosphorylation levels increased from 0 to 15 pSer. Increasing the phosphorylation levels, the pCTD chains exhibited a preference for interacting with HRD over CTD. This observation is reinforced by the decrease in the coordination number of CTD monomers surrounding pCTD monomers, which becomes more pronounced as the chains are more heavily phosphorylated (Fig 5G).

In the dense phase, incrementing phosphorylation levels of pCTD chains in the dense phase above 15 pSer led to $R_G$ values reaching a constant value of 3.6 nm, reflecting the enhanced interaction strength of pCTD with HRD and the pCTD chains are now predominately surrounded by HRD chains. Interestingly, a reduction in anisotropy is also observed with increasing phosphorylation levels, following a prior gradual increase(Fig S11C). This behavior could be attributed to the increased interaction between pCTD and HRD effectively neutralizing the negative charge within pCTD and resulting in more globular conformation of pCTD chains. At this stage, the number of coordinated interactions between CTD and pCTD reached a minimum. This indicates how pCTD's preference for HRD changes as phosphorylation increases. Eventually, this leads to the formation of two distinct phases. The trends observed in the behaviour of $R_G$ and anisotropy further emphasize the shift from a mixed-phase to a segregated phase, comprising pCTD-HRD and CTD, as phosphorylation levels increase, which is in line with the analysis of the density profiles and the evaluation of the size of the largest CTD cluster.

#### Chain conformations in the condensates

Both simulations in large cubic boxes and simulation in extended boxes (slab geometries) showed that the conformation of pCTD is modulated by the pCTD-HRD and CTD phases

(Fig S19). As in the simulation with a large cubic box, pCTD is more extended when it is in the CTD phase than in the pCTD-HRD phase, as tracked by the end-to-end distance of the pCTD chains and the $R_G$.

To explore this further, we calculated the angle $\theta$ which is defined as the angle formed between the eigenvector associated with the largest eigenvalue of chains with the radius vector from the origin at the centre of the condensate (Fig S20). $\cos^2\theta = 1/3$ corresponds to random orientation of the chains. The CTD chains are randomly orientated in the CTD phase (Fig S20A), with $\cos^2\theta$ close to 1/3, which is the value expected for random orientation. At the centre of the pCTD-HRD phase, CTD chains are randomly aligned with $\cos^2\theta = 1/3$( Fig S20B), with a small deviation from random alignment at the interface. On average most pCTD chains in the pCTD-HRD phase are randomly aligned with the $\cos^2\theta$ close to 1/3. Although there is a slight increment in $\cos^2\theta$ observed for chains within the CTD phase, corresponding to a higher $R_G$ and $G_{rr}$, it indicates a tendency for the chains to align perpendicular to the interface. At the interface with the core and shell, there is a decrease in the value $\cos^2\theta$, suggesting a shift towards parallel alignment with the interface. Within the shell region, $\cos^2\theta$ increases to 1/3, implying a more random alignment. Conversely, at the interface of pCTD-HRD with the dilute phase (Fig S20B), $\cos^2\theta$ experiences a significant decrease, signifying a parallel alignment with the interface in this context.

### The conformations of the CTD and pCTD chains are modulated by the two condensates phases

The CTD and pCTD adopt distinct structural conformations depending on the local environment and phase boundaries inside the condensate. Recent theoretical and experimental advances show the profound impact of condensates on the global conformations of proteins within them[12,13], including their orientations and extensions. The conformation of a protein chain within a phase can be understood with the concept of solvent quality from polymer science[14,15].

To understand the conformation of chains in the condensate, we computed parallel and perpendicular components of the gyration tensor.

The components $\langle G_{\parallel}\rangle$ are aligned parallel to the interface, while $\langle G_{\perp}\rangle$ represents the component that is oriented perpendicular to the interface. In the centre of the CTD phase, the extended CTD chains (Fig S12A) are neither aligned parallel nor perpendicular to the condensate interface, with the components of the radius of gyration tensor,[12] $\langle G_{\perp} \rangle$, $\langle G_{\parallel} \rangle$ adopting similar values (Fig S12B).

At the interface of the CTD phase, on average, CTD chains align parallel to the interface, with $\langle G_{\perp} \rangle$ smaller than $\langle G_{\parallel} \rangle$ (Fig S12A, S20A)[16].

Occasionally, some CTD chains visit the pCTD-HRD phase but their $R_G$ is barely affected, even though the phase is enriched in positive and negative charges and thus CTD experiences a reduction in solvent quality. Close to the interface of pCTD-HRD with the dilute aqueous phase the CTD chains collapse as they preferentially interact with the pCTD-HRD phase rather than with the dilute phase.

The conformation of the phosphorylated CTD (pCTD) is modulated differently by the CTD and pCTD-HRD phases. In dilute solution, the degree of phosphorylation greatly influences the global conformation of pCTD. While CTD experiences minimal conformational changes upon transitioning from its parent phase to the pCTD-HRD phase, pCTD undergoes further expansion upon leaving its parent phase and entering the CTD phase. When pCTD is within the CTD phase, it extends even further, with $\langle G_{\parallel} \rangle$ being reduced compared to $\langle G_{\perp} \rangle$, indicating a slight preference for alignment perpendicular to the interface. This preferential alignment perpendicular to the interface enables pCTD to maximize its interactions with positively charged HRD residues in the core. pCTD chains are squeezed at the interface with the dilute phase and $\langle G_{\parallel} \rangle$ are larger than $\langle G_{\perp} \rangle$ as the chains align parallel to the interface. We conclude that the two phases can significantly affect chain conformation due to their different chemical environments and this suggests that the two condensate phases could recruit molecules differentially and underpin specific regulation.

Table S1: IDR sequences of the proteins

| **Protein** | **Sequence** |
|---|---|
| *Homo sapiens* CTD | YSPTSPAYEPRSPGGYTPQSPSYSPTSPSYSPTSPSYSPTSPNSPT<br>SPSYSPTSPSYSPTSPSYSPTSPSYSPTSPSYSPTSPSYSPTSPSY<br>SPTSPSYSPTSPSYSPTSPSYSPTSPSYSPTSPSYSPTSPSYSPTS<br>PSYSPTSPSYSPTSPNYSPTSPNYTPTSPSYSPTSPSYSPTSPNYT<br>PTSPNYSPTSPSYSPTSPSYSPTSPSYSPSSPRYTPQSPTYTPSSP<br>SYSPSSPSYSPASPKYTPTSPSYSPSSPEYTPTSPKYSPTSPKYSP<br>TSPKYSPTSPTYSPTTPKYSPTSPTYSPTSPVYTPTSPKYSPTS<br>PTYSPTSPKYSPTSPTYSPTSPKGSTYSPTSPGYSPTSPTYSLT<br>SPAISPDDSDEEN |
| Histidine Rich Domains (HRD), P-TEFb Cyclin T1[17] | MRIKVHAAADKHNSVEDSVTKSREHKEKHKTHPSNHHHHHN<br>HHSHK HSHSQLPVGTGNKRPGDPKHSSQ |
| *Drosophila melanogaster* CTD | YSPTSPNYTASSPGGASPNYSPSSPNYSPTSPLYASPRYASTTP<br>NFNPQSTGYSPSSSGYSPTSPVYSPTVQFQSSPSFAGSGS<br>NIYSPGNAYSPSSSNYSPNSPSYSPTSPSYSPSSPSYSPTS<br>PCYSPTSPSYSPTSPNYTPVTPSYSPTSPNYSASPQYSPAS<br>PAYSQTGVKYSPTSPTYSPPSPSYDGSPGSPQYTPGSPQYS<br>PASPKYSPTSPLYSPSSPQHSPSNQYSPTGSTYSATSPRYS<br>PNMSIYSPSSTKYSPTSPTYTPTARNYSPTSPMYSPTAPSH<br>YSPTSPAYSPSSPTFEESED |
| *Saccharomyces cerevisiae* CTD | NACFSPTSPTYSPTSPAYSPTSPSYSPTSPSYSPTSPSYSPT<br>SPSYSPTSPSYSPTSPSYSPTSPSYSPTSPSYSPTSPSYSPT<br>SPSYSPTSPSYSPTSPSSPTSPSYSPTSPSYSPTSPAYSPT<br>SPSYSPTSPSYSPTSPSYSPTSPSYSPTSPNYSPTSPSYSP<br>TSPGYSPGSPASSPKQDEQKHNENENSR |

Continued on next page

Table S1: IDR sequences of the proteins (Continued)

| Protein | Sequence |
|---|---|
| FUS low complexity domain (LCD) | NACFSPTSPTYSPTSPAYSPTSPSYSPTSPSYSPTSPSYSPTSP SYSPTSPSYSPTSPSYSPTSPSYSPTSPSYSPTSPSYSPTSPSY SPTSPSYSPTSPSYSPTSPSYSPTSPSYSPTSPAYSPTSPSYSPT SPSYSPTSPSYSPTSPSYSPTSPNYSPTSPSYSPTSPGYSPGS PASSPKQDEQKHNENENSR |
| Ideal CTD 140 mer | YSPTSPSYSPTSPSYSPTSPSYSPTSPSYSPTSPSYSPTSPS YSPTSPSYSPTSPSYSPTSPSYSPTSPSYSPTSPSYSPTSPS YSPTSPSYSPTSPSYSPTSPSYSPTSPSYSPTSPSYSPTSPS YSPTSPSYSPTSPS |

Table S2: Summary of Molecular Dynamics Simulations

| Model | Force Field | Number of Protein Chains | Box Dimension | Time |
|---|---|---|---|---|
| Atomistic | Amber99sb-star-ildn-q | 2 CTD (21mer) | 6.5×6.5×6.5 $nm^3$ | 39.8 $\mu$s |
| Coarse Grained | HPS (300 K) | 2 CTD (21mer) | 7.8×7.8×7.8 $nm^3$ | 10 $\mu$s |
| Coarse Grained | HPS-T (250 K, 280 K, 300 K, 320 K, 340 K, 360 K) | 300 CTD | 15×15×200 $nm^3$ | 5 $\mu$s |
| Coarse Grained | HPS | 100 CTD, 50 pCTD | 15×15×150 $nm^3$ | 10 $\mu$s |

Continued on next page

Table S2: Summary of Molecular Dynamics Simulations (Continued)

| **Model** | **Force Field** | **Number of Protein Chains** | **Box Dimension** | **Time** |
|---|---|---|---|---|
| Coarse Grained | HPS | 100 HRD, Vary pCTD (5,10,20,25,30,50, 70,80,100,110,150, 200,250) | 15×15×150 $nm^3$ | 12 $\mu$s |
| Coarse Grained | HPS | 100 CTD, 100 pCTD, 250 HRD | 15×15×100 $nm^3$ | 15 $\mu$s |
| Coarse Grained | HPS | 600 CTD | 100×100×100 $nm^3$ | 10 $\mu$s |
| Coarse Grained | HPS | 100 CTD, 200 FUS | 15×15×200 $nm^3$ | 5 $\mu$s |
| Coarse Grained | HPS | 100 pCTD, 200 FUS | 15×15×200 $nm^3$ | 5 $\mu$s |
| Coarse Grained | HPS | 200 CTD, 200 pCTD, 400 HRD, 100 FUS | 200×200×200 $nm^3$ | 5 $\mu$s |
| Coarse Grained | HPS | 300 CTD, Vary pCTD (30,50,100,150,200,), Vary HRD (30,50,100,150,200,) | 200×200×200 $nm^3$ | 6 $\mu$s |

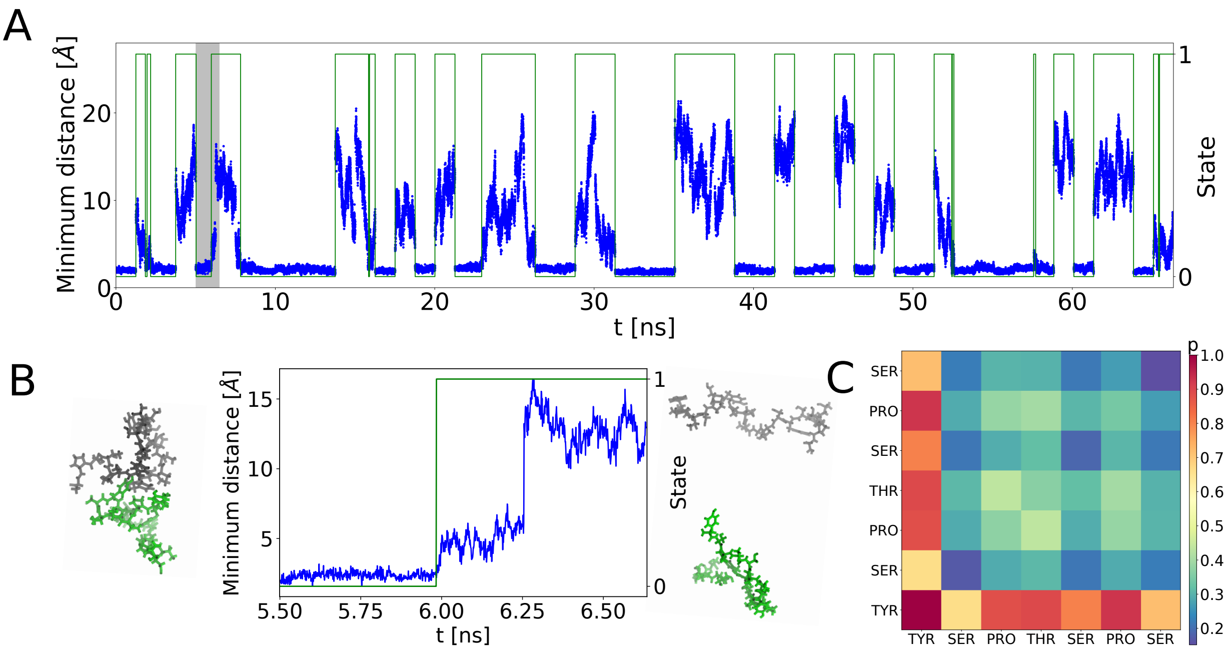


Figure S1. Atomistic molecular dynamics simulations of CTD fragments. A) The association and dissociation dynamics of CTD fragments is tracked by computing the minimum distance between two fragments in this 65 ns segment of a 129 ns trajectory. The system transitions between bound (0) and unbound states (1). The highlighted area is enlarged in B) B) Association and dissociation of CTD fragments corresponding to the movie S1. Snapshots illustrate the bound and unbound states of two CTD chains, with one chain shown in grey and the other in green respectively. Water and ions are omitted for clarity C) Contact map for two all-atom CTD chains showing Tyr-Tyr and Tyr-Pro interactions. A pair of residues is in contact if any atom is below a threshold of $2^{1/6}(\sigma_i + \sigma_j)/2$, where $\sigma$ is the diameter of the residues. The simulation is 39 µs long.

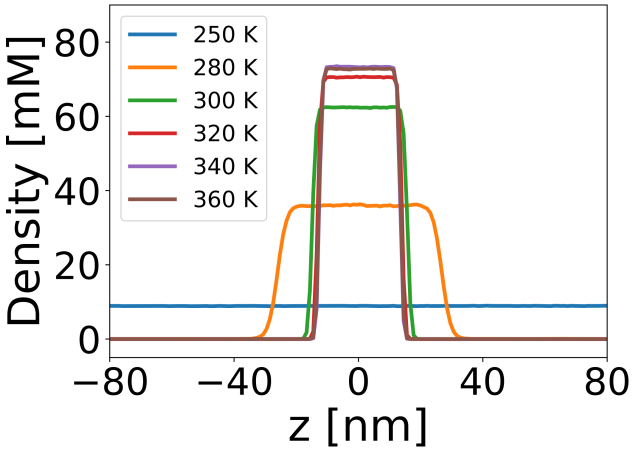


Figure S2. Temperature-Dependent Phase Behavior of C. elegans CTD. The density profiles of condensates at different temperatures. The system phase separates at higher temperatures.

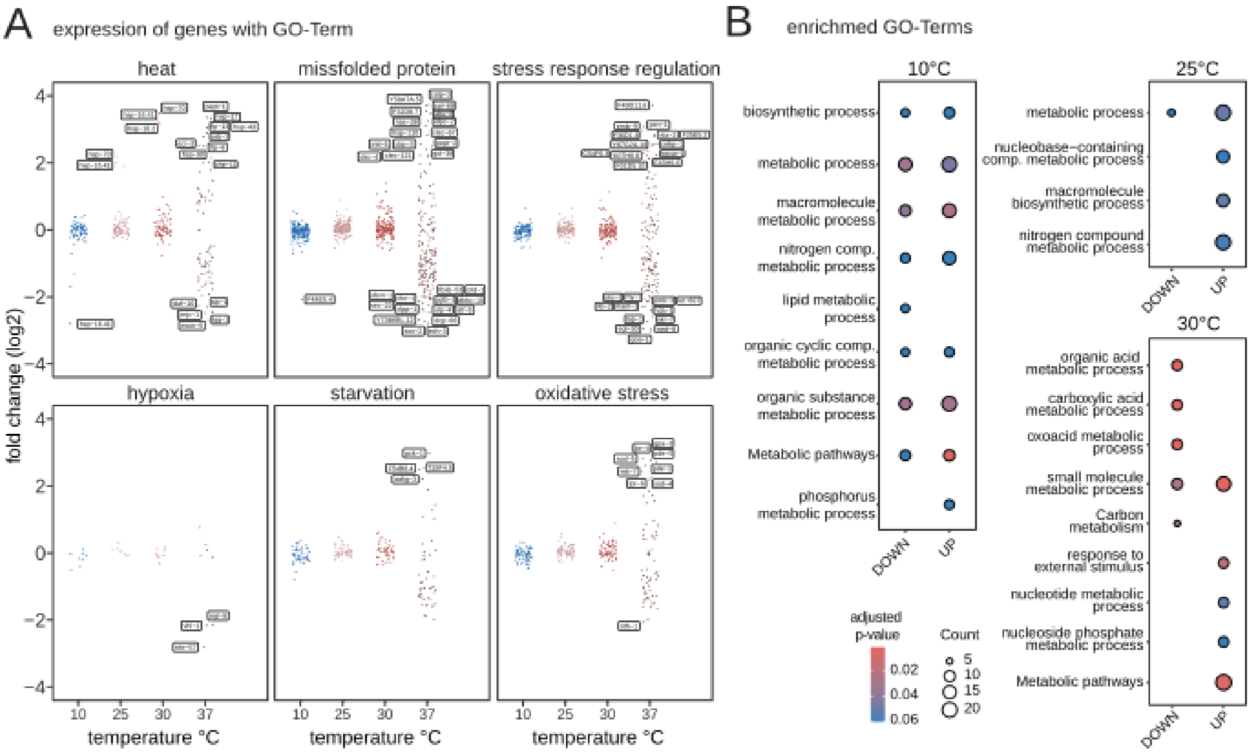


Figure S3. A) Expression changes as mean fold change (log2) over 20°C for genes with the indicated GO-Terms involved in a range of stress responses. Genes with a mean fold change $> 2$ or $< -2$ are labelled. Gene expression at 10°C, 25°C and 30°C are compared to expression changes at heat stress (37°C)[18]. B) GO-Term and KEGG pathway analysis of UP and downregulated genes.

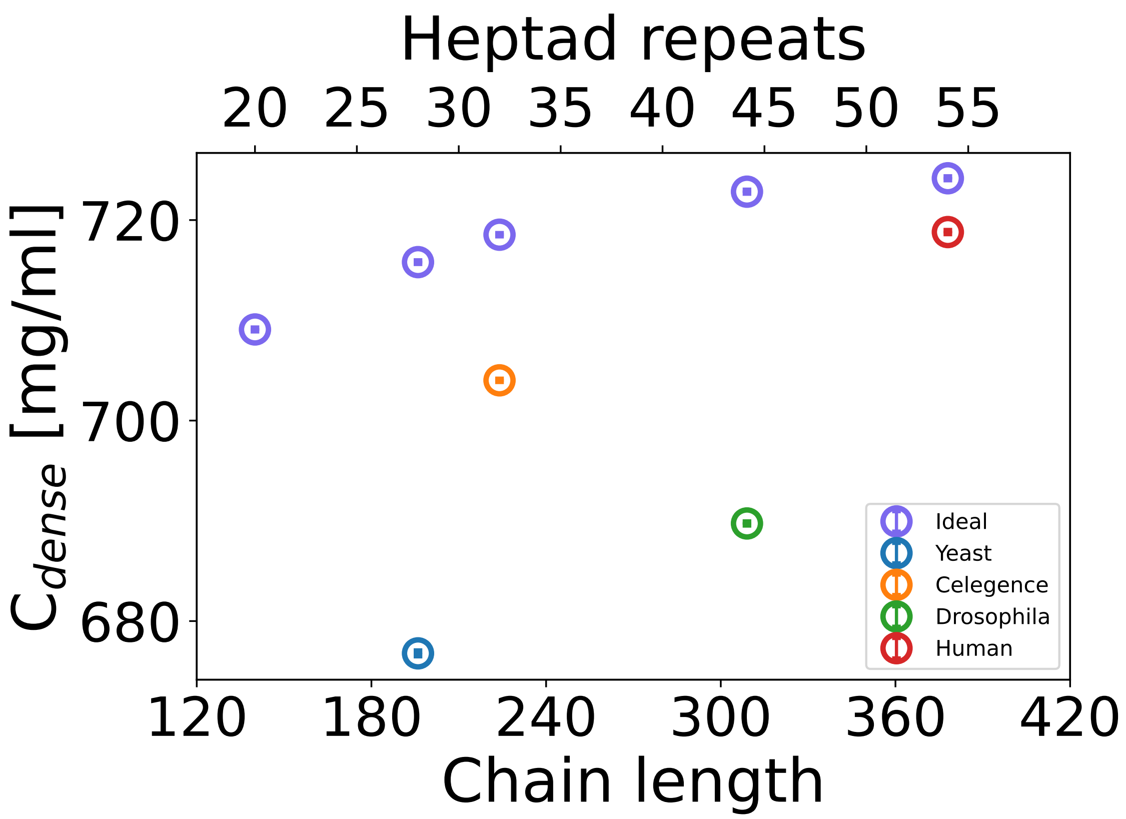


Figure S4. A) Comparison of the dense concentration of real and ideal sequences from *Drosophilia melanogaster*, *Saccharomyces cerevisiae*, and *Homo sapiens*. The ideal sequences are less prone to phase separation compared to the real sequence. Shorter CTD sequences exhibited reduced phase separation compared to longer ones.

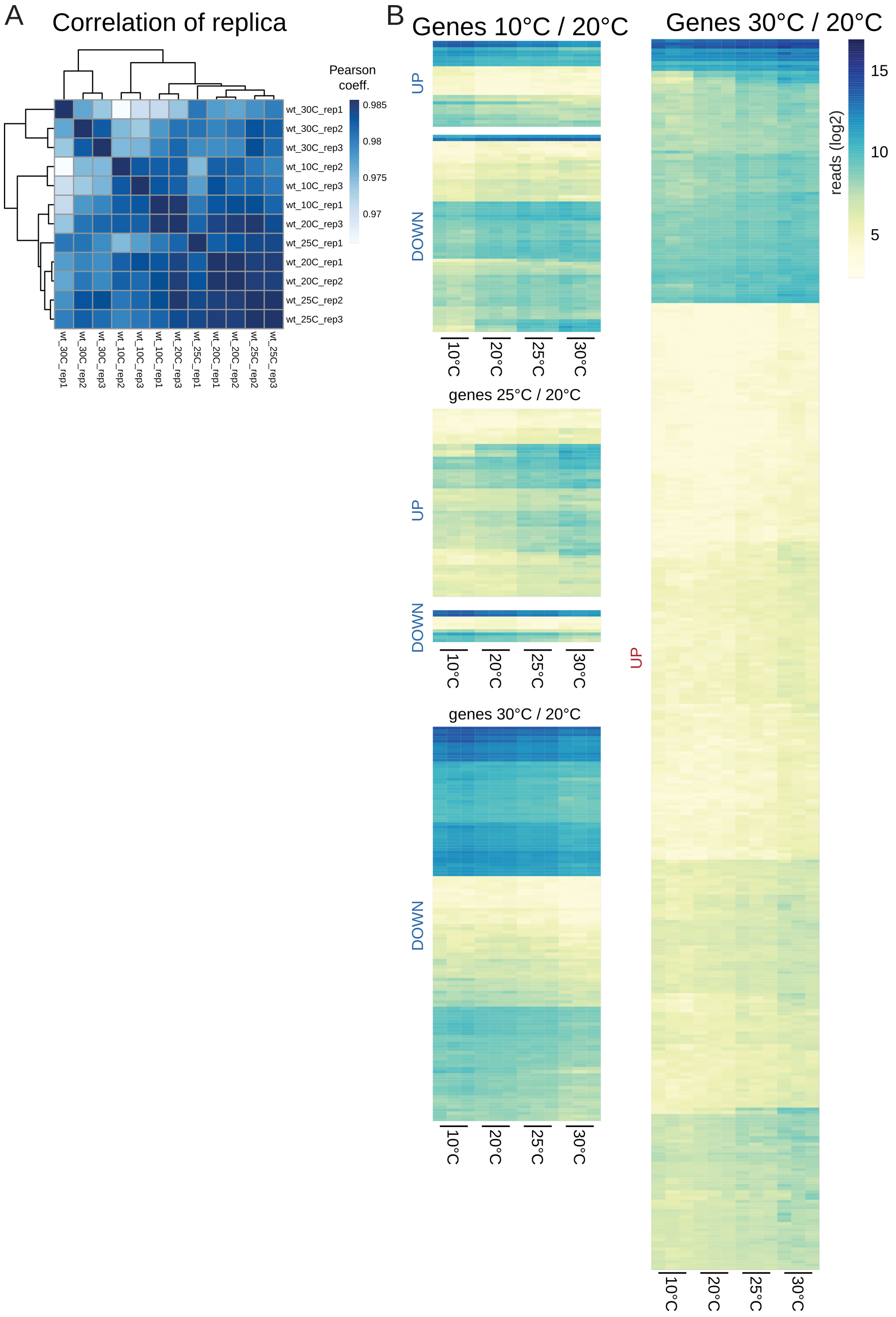


Figure S5. (Caption next page.)

Figure S5. (Previous page.)Exposing *C. elegans* embryos to a range of temperatures from 10°C-30°C results in a gradual transcriptional response. A) Heatmap showing the Pearson correlation coefficient calculated for each combination of replica and temperature of the RNA-seq B) Heat maps showing the gene expression level as normalized reads per gene (log2) of embryos cultured at 20°C (standard laboratory conditions) and embryos exposed to 10°C, 25°C, or 30°C for 2hrs. (RNA-seq) of significantly changed genes (FDR $< 0.05$), red significantly upregulated genes (fold change $> 2$), and blue significantly downregulated genes (fold change $< -2$). See also Fig 3A.

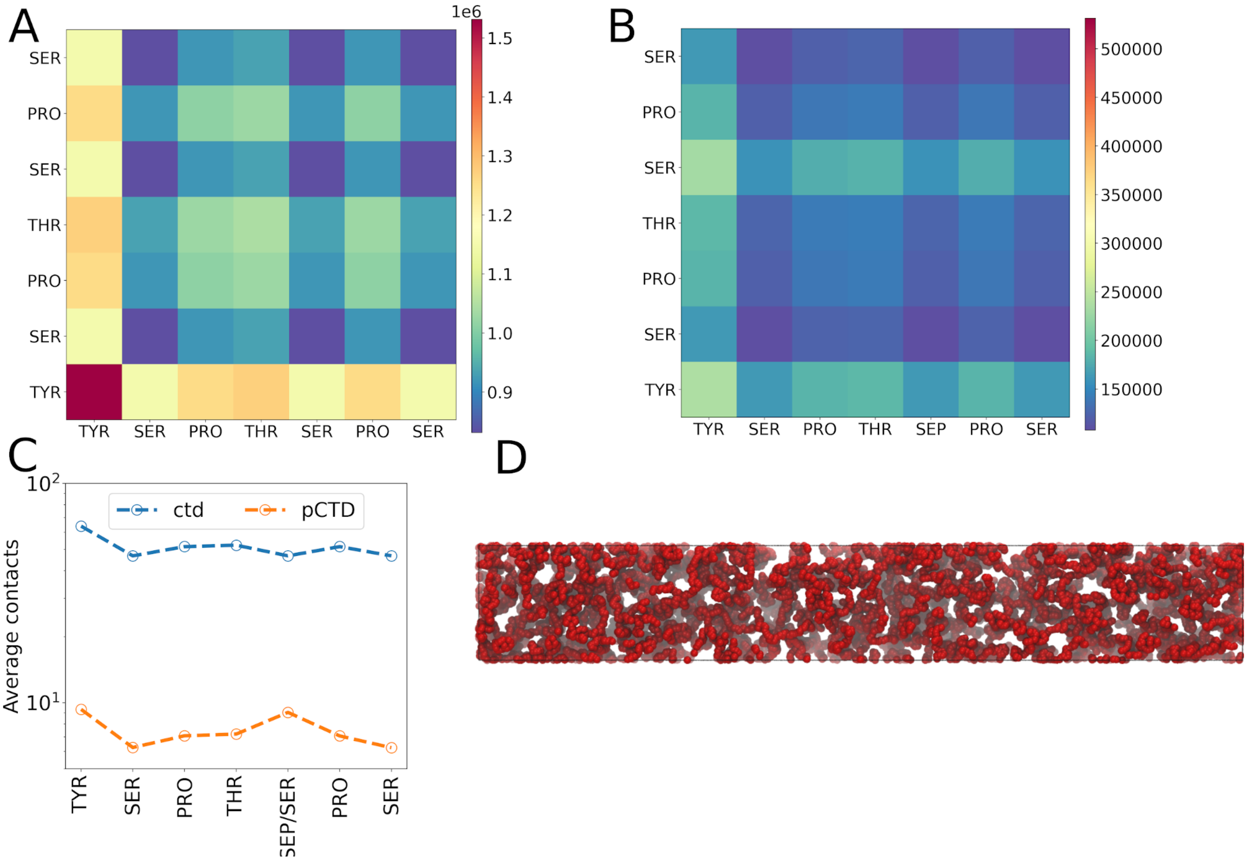


Figure S6. (A) Contact maps depict intermolecular interactions among residues within the CTD chains (B) Between the CTD and phosphorylated CTD (pCTD) chains with all the $Ser_5$ phosphorylated.(C) The average residue contacts between the CTD and pCTD chains. The strength of CTD-CTD interactions is approximately tenfold higher than that of pCTD-CTD interactions as judged by the number of contacts, highlighting a substantial difference in intermolecular affinities. D) Phosphorylation of all $Ser_5$ positions of pCTD (red) dissolves the condensate.

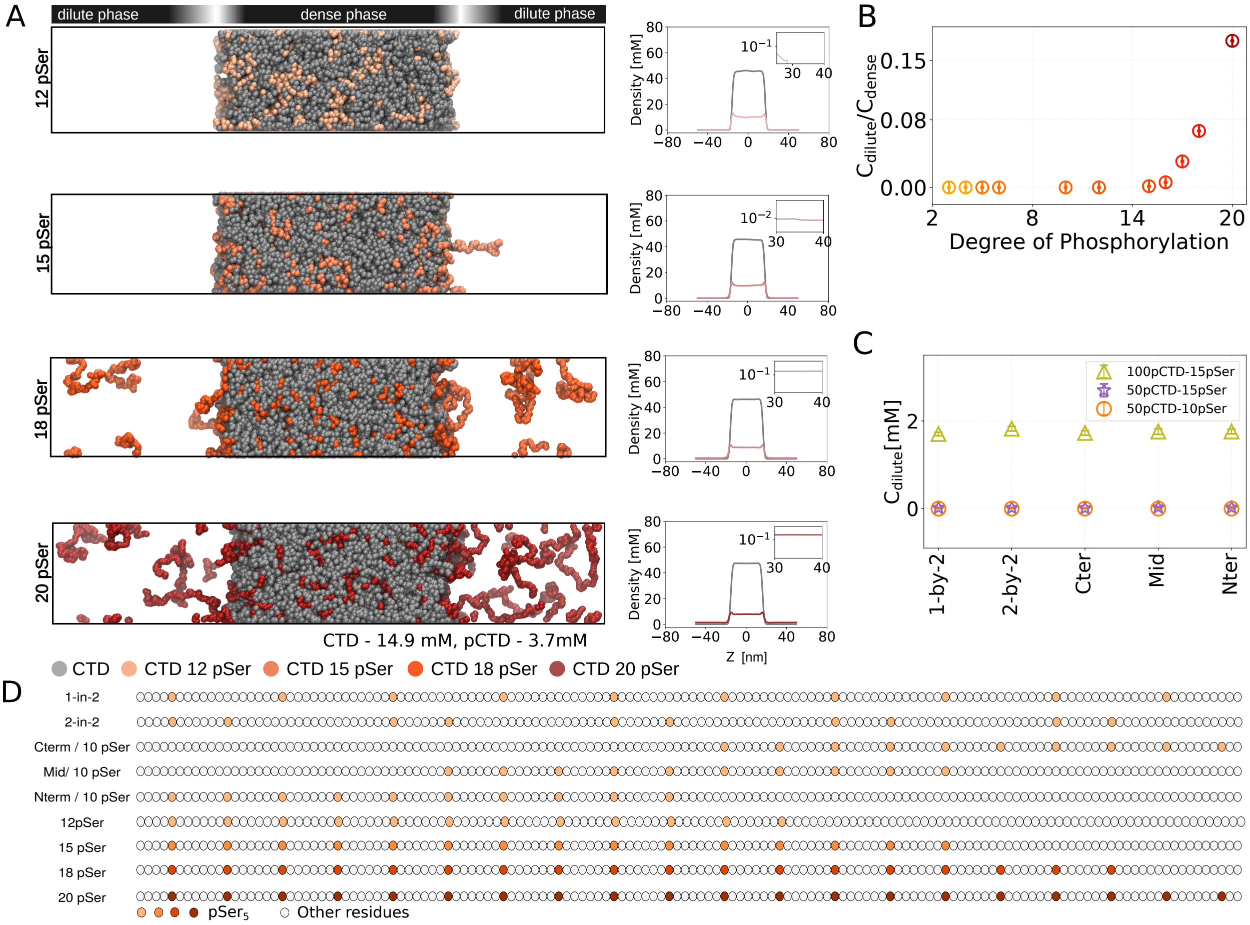


Figure S7. Phase behavior of 100 CTD and 50 pCTD chains. Visualization of CTD and pCTD in molecular dynamics simulations in slab geometry of box dimension 15 nm×15 nm×100 nm. A) The degree of phosphorylation of pCTD chains varies in each simulation. Density plots for simulation with CTD and pCTD with 10pSer, 12pSer, 18pSer, and 20pSer respectively. B) Quantifying $c_{dilute}/c_{dense}$ of partially phosphorylated pCTD chains as the $Ser_5$ phosphorylation increases. C) Variation in $c_{dilute}$ of pCTD with pSer residues distributed in five distinct patterns. D) Distribution of $pSer_5$ residues in the 140mer pCTD.

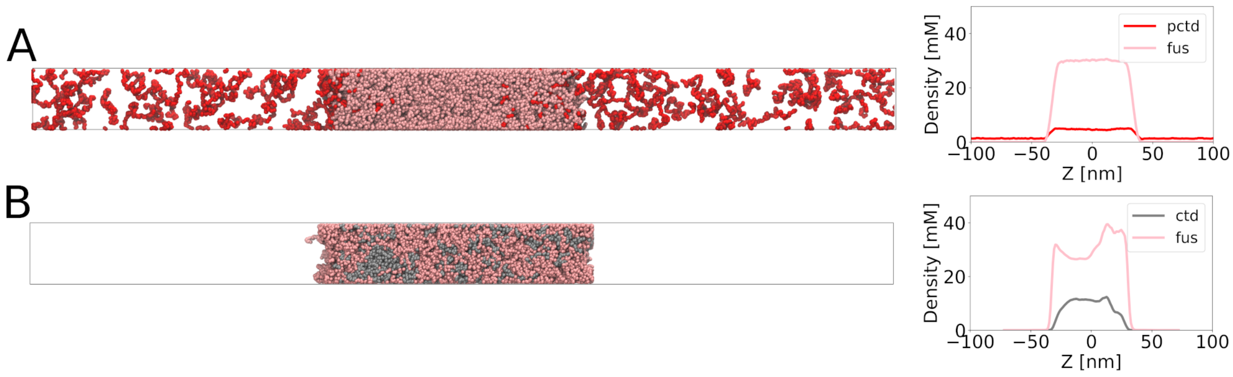


Figure S8. FUS (pink) interaction with A) pCTD (red) and B) CTD (grey). FUS exhibits stronger interaction with CTD compared to pCTD. Recruitment of CTD to the FUS condensates is evident for unphosphorylated CTD, whereas the interaction strength is reduced with pCTD.

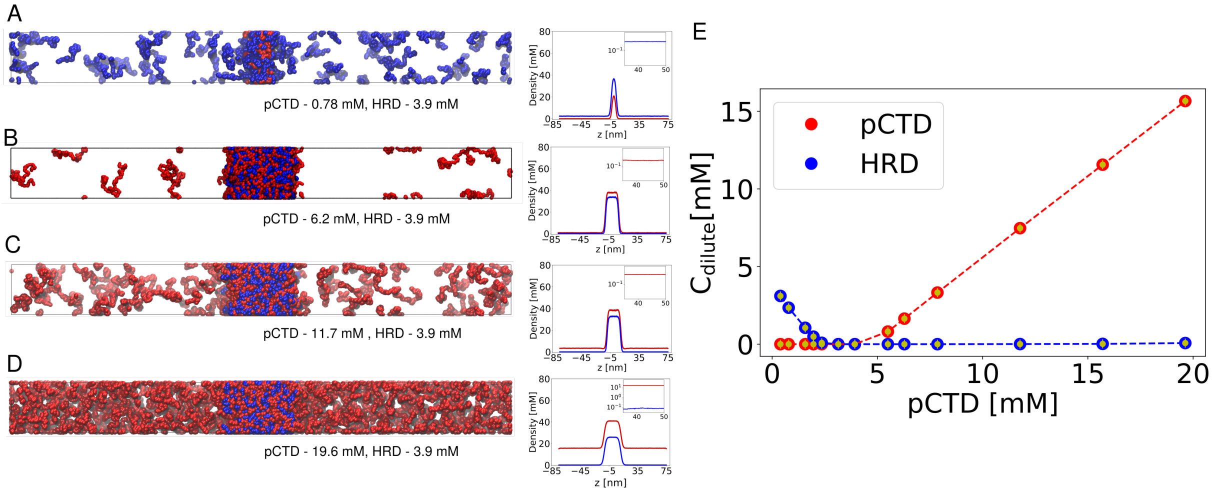


Figure S9. Co-phase separation of HRD with CTD and pCTD in slab simulations A-D) Simulation snapshots depicting the effect of increasing pCTD concentration while maintaining a constant HRD concentration in a slab simulation geometry. Density profiles for pCTD and HRD correspond to the simulations shown on the right side. E) Variation of dilute phase concentrations of pCTD and HRD as the overall pCTD concentration increases.

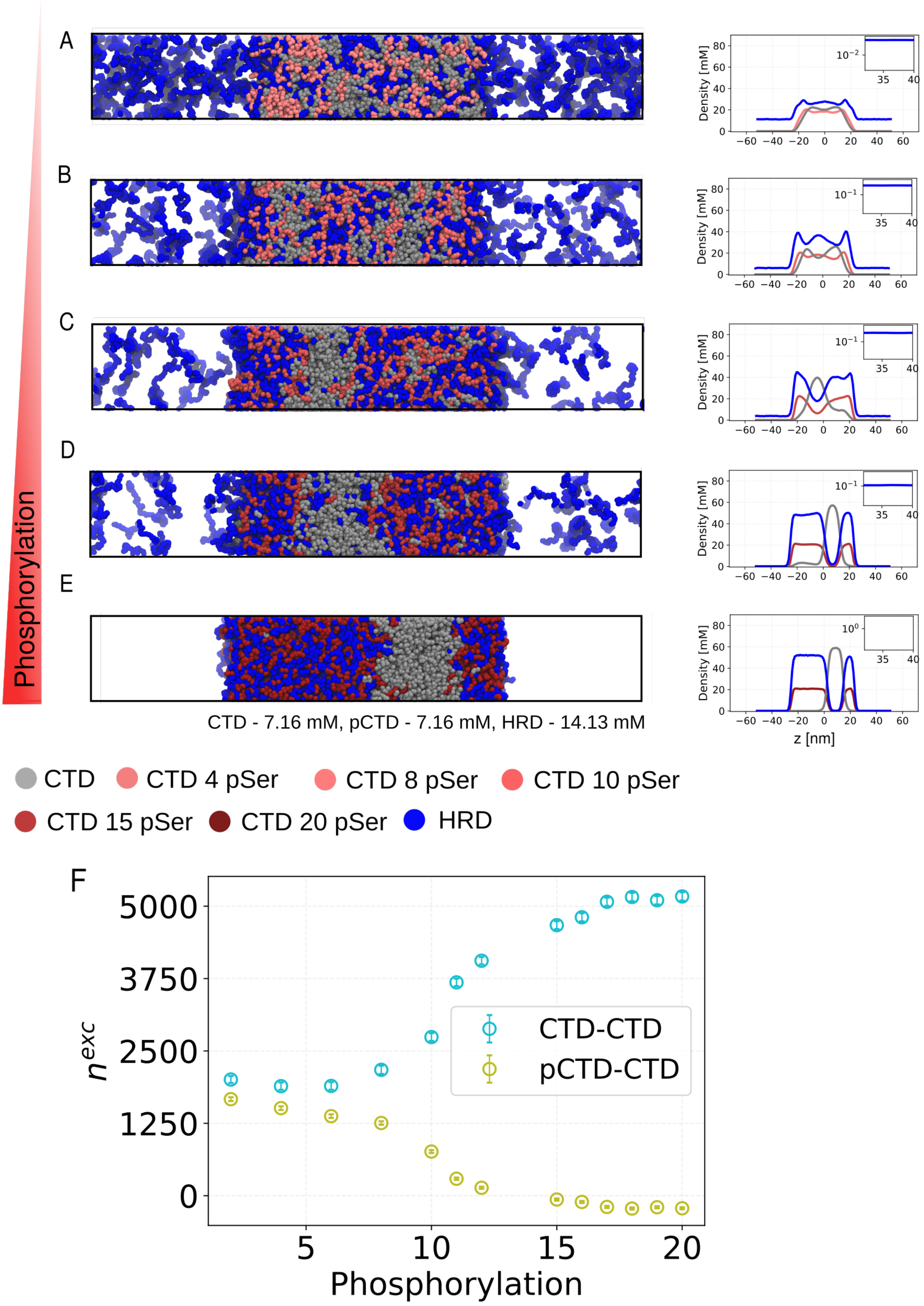


Figure S10. (Caption next page.)

Figure S10. (Previous page.) Mixing and de-mixing of the dense phase of 100 CTD, 100 pCTD, and 250 HRD chains, with increasing the degree of pCTD phosphorylation. A) Snapshot and density profile for the simulations where 4pSer B) 8pSer C) 10pSer D) 15pSer E) 20 pSer of pCTD are phosphorylated. G) Comparison of the excess of CTD monomers surrounding a phosphorylated pCTD (CTD-pCTD) and surrounding another CTD (CTD-CTD), with increasing phosphorylation levels, denoted as $n^{exc}$. This metric quantifies the number of protein-protein contacts and is calculated using Equation 12.

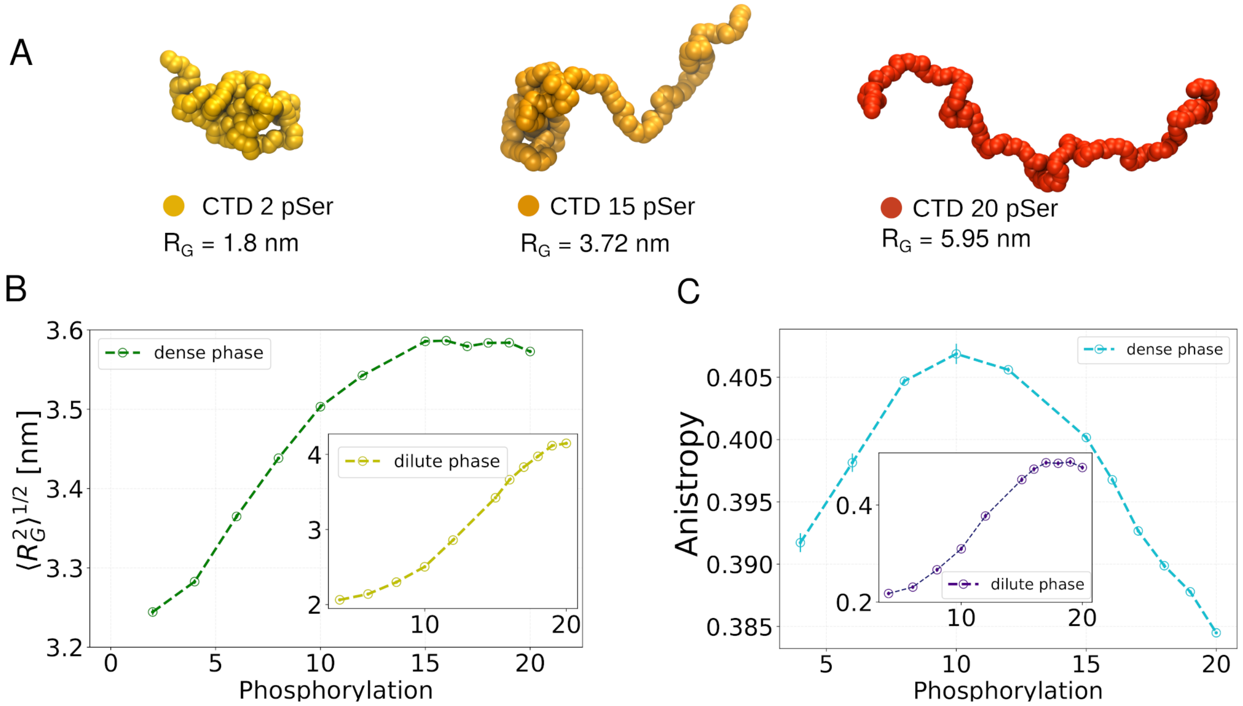


Figure S11. Variation in the pCTD conformation with the degree of phosphorylation in the slab simulations with CTD, pCTD, and HRD (refer to Figure 5). $R_G$ and Anisotropy of pCTD in dense and dilute phase. A) The conformations of pCTD chains in the dilute phase with an increase in the number of phosphorylated $Ser_5$ B) Variation in $R_G$ of pCTD chains in the dense phase with the degree of phosphorylation, with an inset showing the variation of $R_g$ in the dilute phase. C) Variation in the anisotropy of pCTD chains with an increase in phosphorylation of pCTD in the dense phase. Iset depicting the variation in the dilute phase.

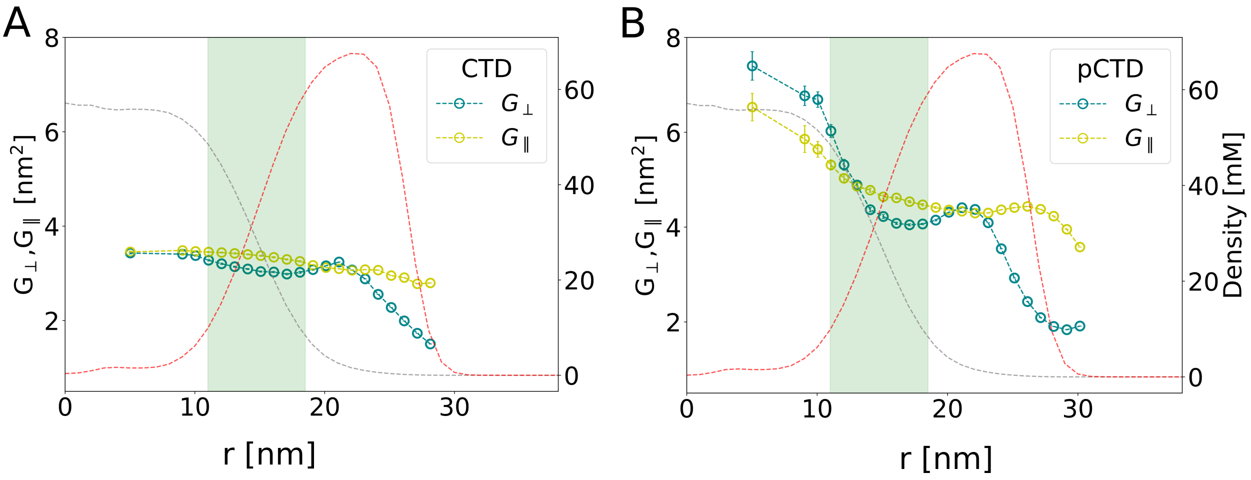


Figure S12. A) Changes in $R_G$ and diagonal components of the gyration tensor for CTD chains, measured from the droplet's centre. B) Variation in $R_G$ and diagonal components of the gyration tensor for pCTD chains, measured from the droplet's centre.

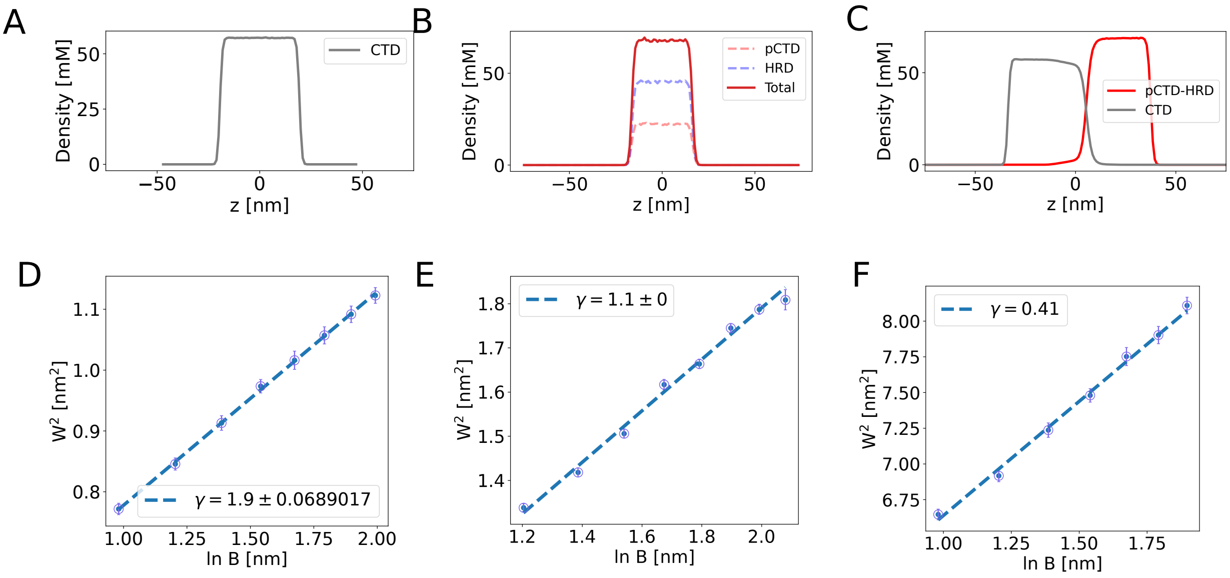


Figure S13. Analysis of interfacial tension from slab simulations using Capillary Wave Theory (CWT) with the block analysis method. Panels A-C display density profiles for CTD, pCTD-HRD, and CTD-pCTD-HRD systems. Panels D-F illustrate the apparent interfacial width $W^2$ with block size B. The interface region is divided into rectangular segments. The interface region is divided into rectangular segments $W_0^2 + \frac{K_B T}{2\pi\gamma} \ln\left(\frac{L}{B}\right)$, from which the interfacial tension is determined.

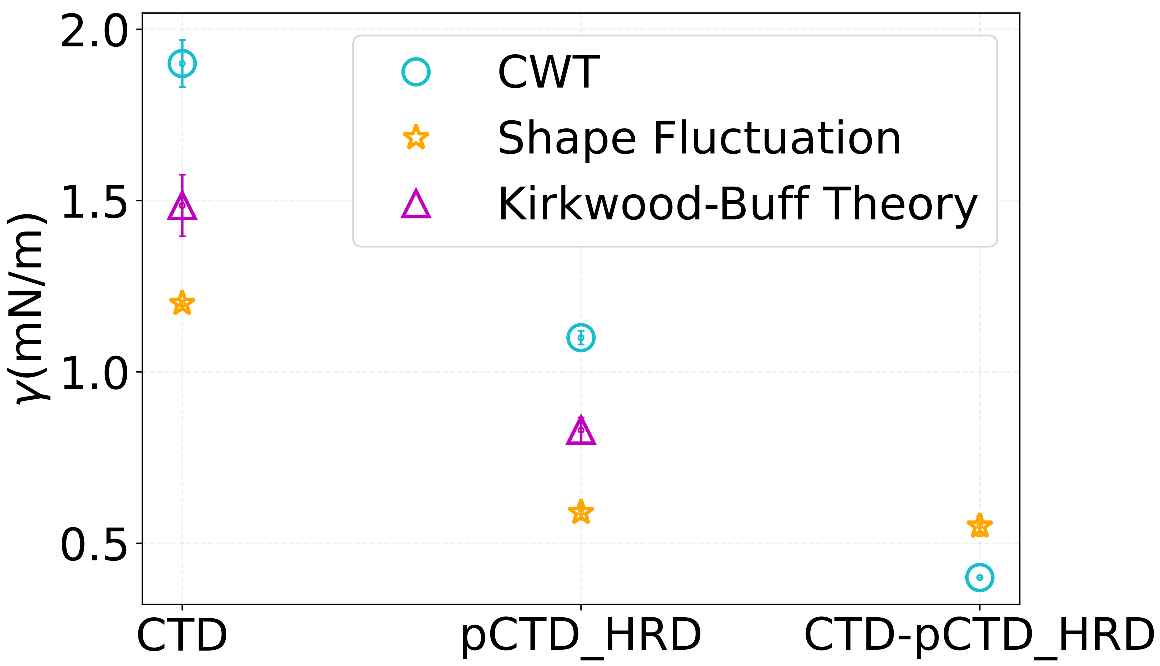


Figure S14. Comparison of interfacial tension obtained from different methods: Capillary Wave Theory (CWT) with interfacial width calculation, Shape Fluctuation of Droplet, and Kirkwood-Buff Theory. Comparison among CTD, pCTD-HRD condensates, and the interface between CTD and pCTD-HRD.

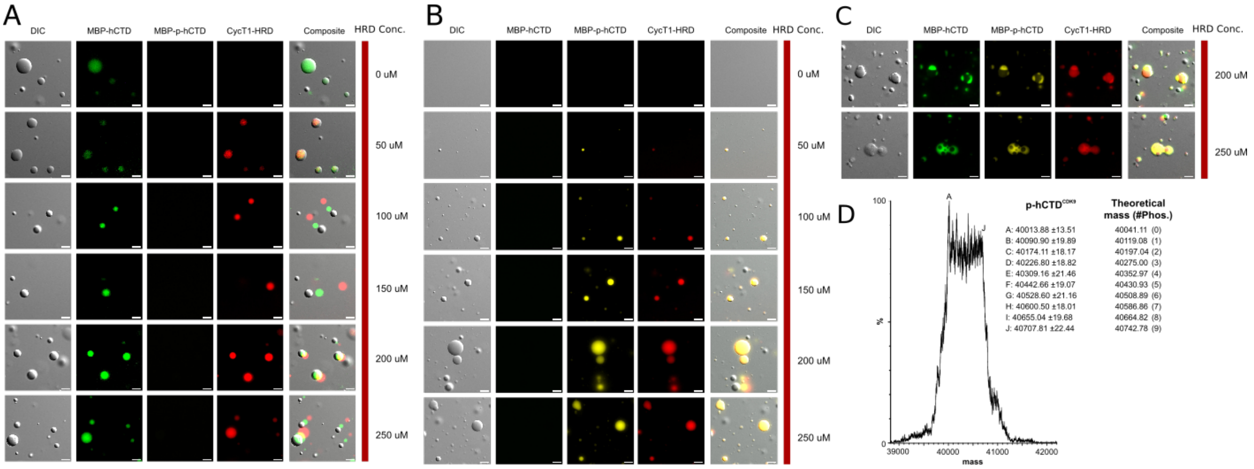


Figure S15. A) HRD co-recruitment into hCTD droplets. Differential interference contrast (DIC) shows MBP-hCTD (AF488; green) and HRD (AF647; red) co-localized in protein condensates. Composite pictures show the different levels of mobility for the droplets. The incremental concentration of HRD is indicated. B) Liquid-liquid phase separation of phosphorylated hCTD assisted by HRD. Differential interference contrast (DIC) shows MBP-p-hCTD (AF594; yellow) and HRD (AF647; red) condensates. HRD concentration is indicated. C) Droplet compartments induced by HRD. Composite pictures show droplets formed by different components in function of HRD's concentration of 200 and 250 M. D) Mass spectrometry for p-hCTD. Masses are detected (p-hCTDCDK9 column) and connected to different levels of phosphorylation of hCTD using CDK9 (Theoretical mass column)

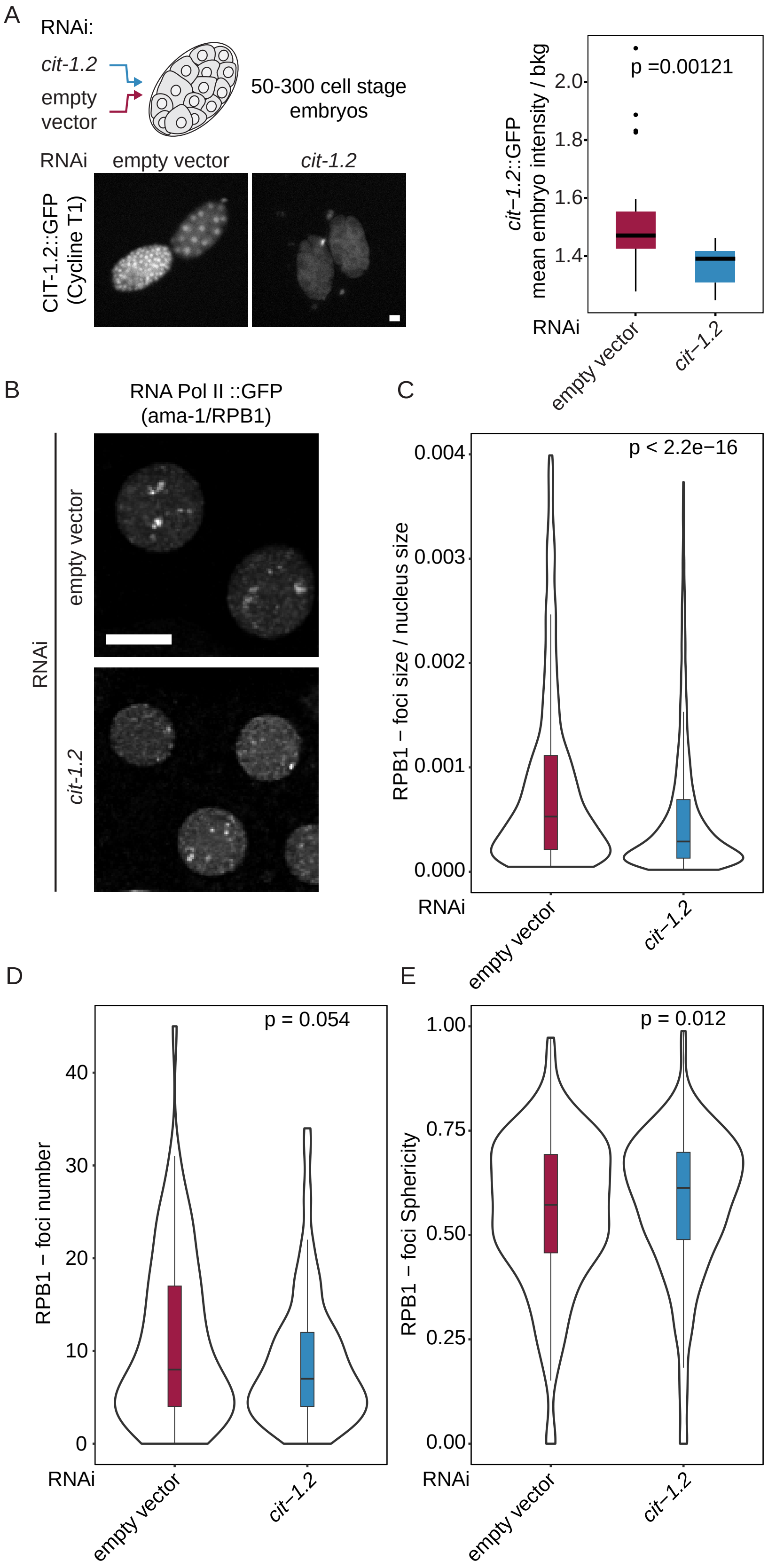


Figure S16. Loss of Cycline T1 results in smaller and rounder RNA Pol II foci in C. elegans embryos (Next page.)

Figure S16. (Previous page.)(A) Control images of embryos expressing the endogenous CIT-1.2 (Cycline T1) fused to GFP treated with RNAi against cit-1.2, or empty vector. Mean GFP intensity was quantified per embryo and normalized to the background. (B) Representative images of RNA Pol II foci in C. elegans embryos treated with empty vector, or Cycline T1 (cit-1.2) RNAi (at 20°C). size bar = 4um (C) The size of foci relative to the total nuclear volume. (D) The number of foci per nucleus. (E) The Sphericity of RNA Pol II foci. P-values (Wilcoxon rank sum test) are indicated above the individual comparisons. N=3, n(empty vector)=147 nuclei, n(cit-1.2)=206

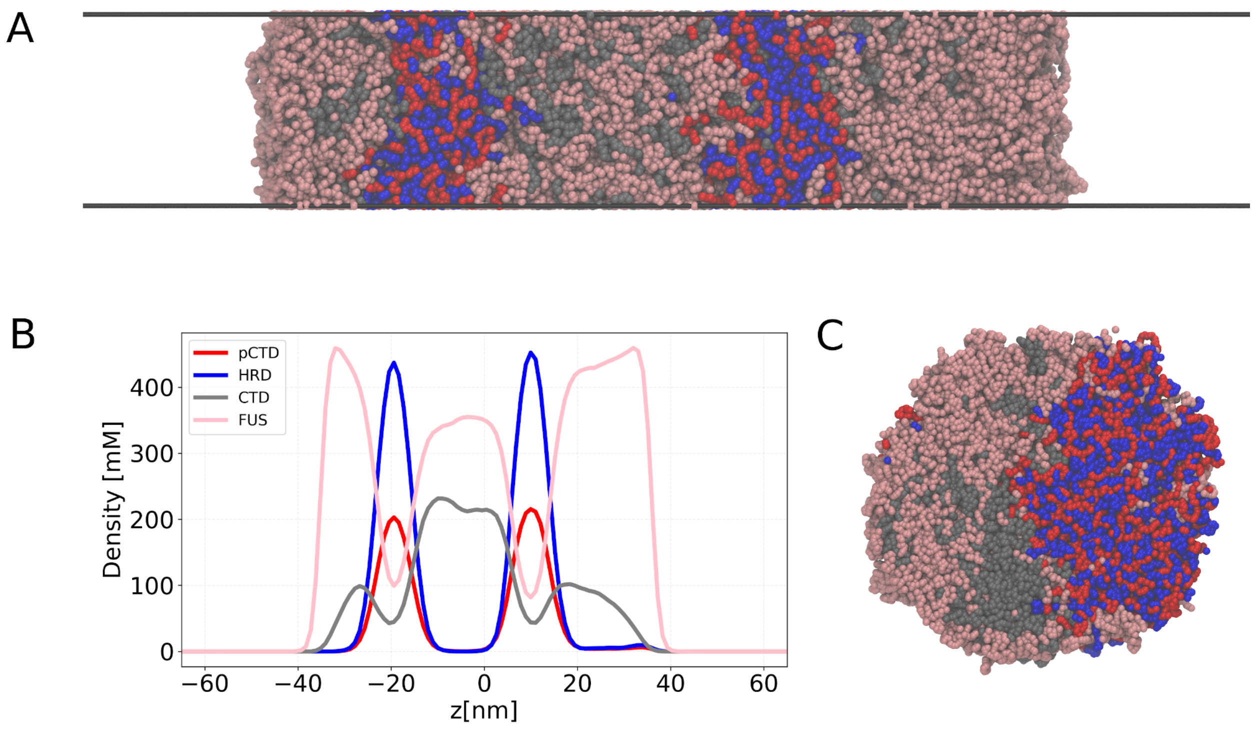


Figure S17. A) Simulation of pCTD, CTD, HRD, and FUS (Pink). B) Density profile showing FUS interactions with CTD. C) Cross section of the droplet. The droplet has partial engulfment morphology.

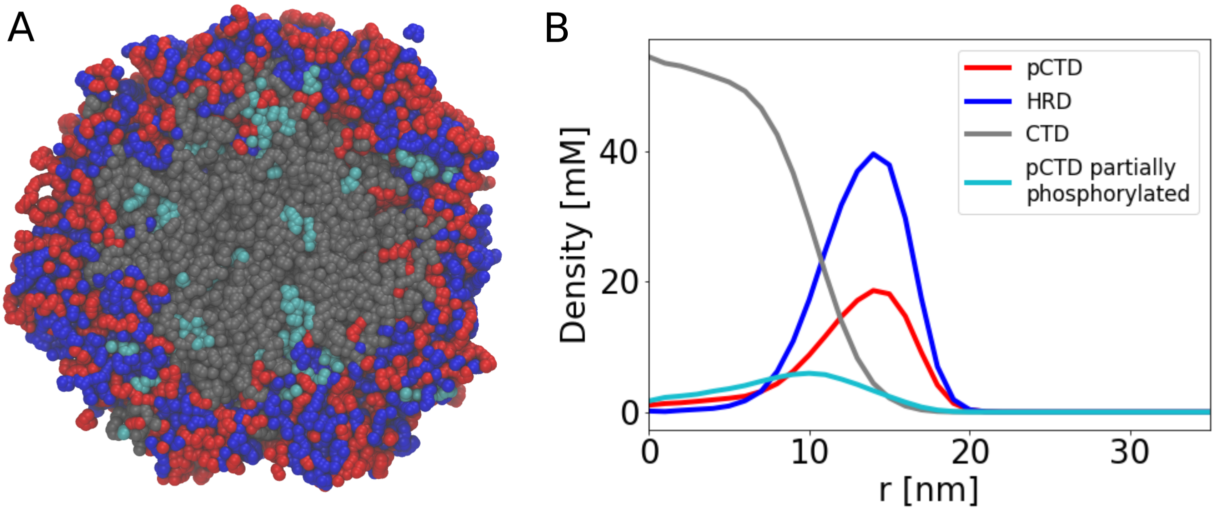


Figure S18. Interactions of partially phosphorylated CTD with the CTD and pCTD-HRD phases. A) Picture shows cut through a droplet of partially phosphorylated (cyan), fully phosphorylated CTD (red), unphosphorylated CTD, and HRD from coarse-grained molecular dynamics. In the partially phosphorylated CTD, $\mathrm{Ser}_5$ within the initial 10 heptad is phosphorylated. B) Density profile of the system.

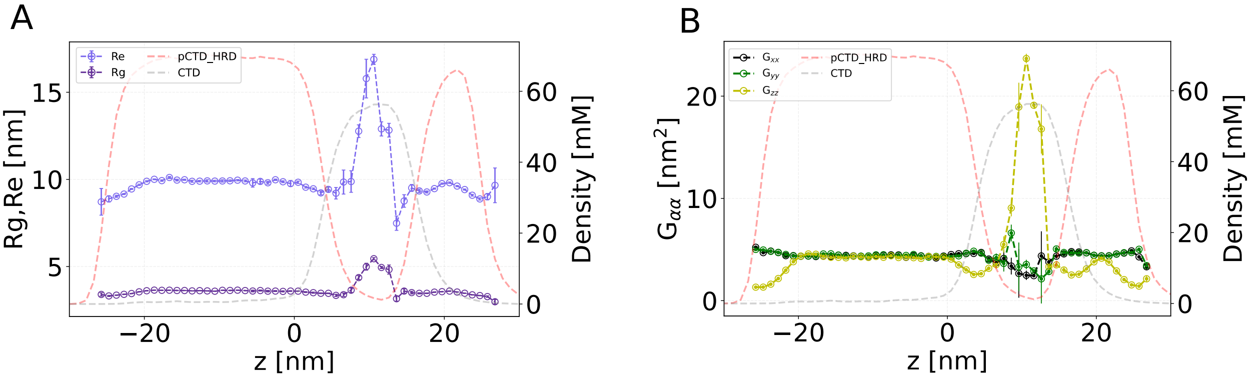


Figure S19. The diagonal elements of the gyration tensor for pCTD chains in the multiphasic condensate in a slab simulation, denoted as $\langle G_{xx} \rangle$, $\langle G_{yy} \rangle$, and $\langle G_{zz} \rangle$, are plotted against the distance from the center of the box. This observed pattern aligns with findings from droplet simulation. Within their assigned phase, pCTD chains exhibit confinement, while in the less favorable CTD phase, they adopt an extended structure. Across all interfaces, the $\langle G_{rr} \rangle$ component decreases, indicating chains try to align parallel to the interface and experience compression.

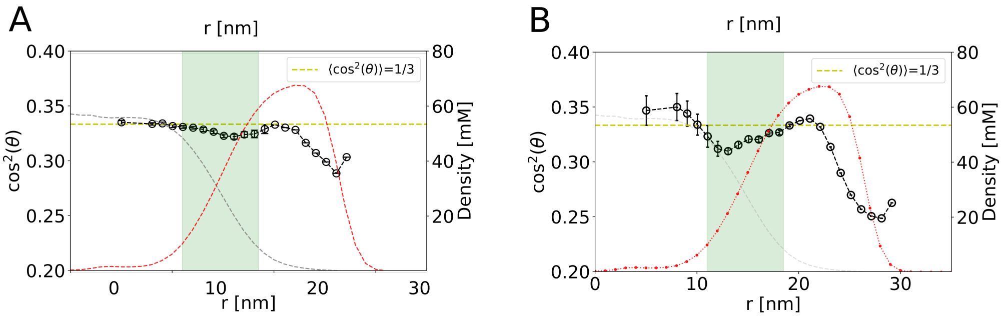


Figure S20. Variations in $\cos^2\theta$ from the droplet's center for A) CTD and B) pCTD chains.
